# Prostate cancer-induced endothelial-to-osteoblast transition generates an immunosuppressive bone tumor microenvironment

**DOI:** 10.1101/2023.11.30.569496

**Authors:** Guoyu Yu, Paul G. Corn, Celia Sze Ling Mak, Xin Liang, Miao Zhang, Patricia Troncoso, Jian H. Song, Song-Chang Lin, Xingzhi Song, Jingjing Liu, Jianhua Zhang, Christopher J. Logothetis, Marite P. Melancon, Theocharis Panaretakis, Guocan Wang, Sue-Hwa Lin

## Abstract

Immune checkpoint therapy has limited efficacy for patients with bone metastatic castrate-resistant prostate cancer (bmCRPC). In this study, we revealed a novel mechanism that may account for the relative resistance of bmCRPC to immune checkpoint therapy. We found that prostate cancer (PCa)-induced bone via endothelial-to-osteoblast (EC-to-OSB) transition causes an ingress of M2-like macrophages, leading to an immunosuppressive bone tumor microenvironment (bone-TME). Analysis of a bmCRPC RNA-seq dataset revealed shorter overall survival in patients with an M2-high versus M2-low signature. Immunohistochemical (IHC) analysis showed CD206^+^ M2-like macrophages were enriched in bmCRPC specimens compared with primary tumors or lymph node metastasis. In osteogenic PCa xenografts, CD206^+^ macrophages were enriched adjacent to tumor-induced bone. FACS analysis showed an increase in CD206^+^ cells in osteogenic tumors compared to non-osteogenic tumors. Genetic or pharmacological inhibition of the EC-to-OSB transition reduced aberrant bone and M2-like macrophages in osteogenic tumors. RNAseq analysis of tumor-associated macrophages from osteogenic (bone-TAMs) versus non-osteogenic (ctrl-TAMs) tumors showed high expression of an M2-like gene signature, canonical and non-canonical Wnt pathways, and a decrease in an M1-like gene signature. Isolated bone-TAMs suppressed T-cell proliferation while ctrl-TAMs did not. Mechanistically, EC-OSB hybrid cells produced paracrine factors, including Wnts, CXCL14 and LOX, which induced M2 polarization and recruited M2-like TAMs to bone-TME. Our study thus links the unique EC-to-OSB transition as an “upstream” event that drives “downstream” immunosuppression in the bone-TME. These studies suggest that therapeutic strategies that inhibit PCa-induced EC-to-OSB transition may reverse immunosuppression to promote immunotherapeutic outcomes in bmCRPC.

**Significance:** The insight that prostate cancer-induced bone generates an immunosuppressive bone tumor microenvironment offers a strategy to improve responses to immunotherapy approaches in patients with bone metastatic castrate-resistant prostate cancer.

## Introduction

Bone is the dominant site of prostate cancer (PCa) metastases and occurs in >70% of patients with castrate-resistant disease (1). Despite the development of cytotoxic agents, second generation androgen-receptor targeting therapies, and radioligand therapies, metastatic progression in bone signifies lethal disease (2). Although immune checkpoint therapy (ICT) has significantly improved overall survival in a number of solid tumors, bone metastatic castrate-resistant prostate cancer (bmCRPC) is relatively unresponsive (3,4). Several mechanisms may contribute to the relative resistance of bmCRPC to ICT. For example, studies by Jiao et al. (3) showed that ICT led to an increase in Th17 rather than Th1 subsets in bone marrow mediated by the release of TGF-β during osteoclast-mediated bone resorption. More recently, Kfoury et al. (5) performed single-cell RNAseq analysis of bone from patients undergoing laminectomy due to spinal cord compression and found that M2-polarized macrophages were enriched in PCa bone metastasis. These observations suggest that M2-like macrophages in bone metastasis may play a role in promoting PCa progression in bone. However, mechanisms that lead to the enrichment of M2-like macrophages in the PCa-bone metastasis remain unknown.

A unique feature of PCa bone metastasis is the induction of aberrant bone overgrowth leading to osteoblastic lesions (6). This unique “bone-forming” feature can be detected clinically using technetium-99m bone scans and elevations in serum alkaline phosphatase levels associated with osteoblast activity. This phenotype is in marked contrast to most other solid tumors, including renal, lung and breast carcinomas, which induce osteolytic bone lesions. We and others have shown that PCa-induced aberrant bone formation supports tumor progression and resistance to therapies (7,8). Several lines of evidence suggest that bone morphogenetic protein (BMPs), including BMP4 (9,10), BMP6 (11), and BMP7(12) secreted from PCa cells are involved in bone forming lesions, consistent with a role of BMPs in promoting osteoblast differentiation. As a result, PCa-induced aberrant bone formation was initially thought to result from the expansion of existing osteoblasts (6). However, studies by Lin et al. (13) demonstrated that tumor cells secrete BMP4, which induces endothelial-to-osteoblast (EC-to-OSB) transition as a mechanism of PCa-mediated bone-forming metastases.

In this study, we sought to test the hypothesis that tumor-induced stromal reprogramming through the EC-to-OSB transition mechanistically contributes to the immunosuppressive bone-TME. To approach this, we used osteogenic vs non-osteogenic tumor models in mice. Our studies reveal a novel mechanism of immunosuppression in the bone-TME due to the ingress of M2 macrophages mediated by EC-OSB transition.

## Results

### High levels of M2-like macrophages in bone metastasis correlate with shorter survival in mCRPC patients

To examine the involvement of M2-like macrophages in PCa progression, we first used an M2-like macrophage gene signature derived from Cassetta et al. (14) to examine the castrate-resistant prostate cancer (CRPC) bone metastasis from a well annotated mCRPC RNA-seq dataset (15). We found that bone metastasis samples could be classified into M2-high and M2-low subsets using hierarchical clustering (**Fig. 1A**). In addition, several immunosuppressive genes, including *CD274* (encodes PD-L1), *PDCD1LG2* (encodes PD-L2), *CSF1*, and *TGFB1,* which are associated with M2-like macrophages, had higher expression in M2-high bone metastasis samples, although the p value for *TGFB1* (0.06) did not reach statistical significance (**Fig. 1B**). We also observed a trend towards shorter survival in patients with the M2-high compared to M2-low signature (logRank P=0.0664) (**Fig. 1C**). Of note, since our studies sought to elucidate how bone-forming metastases modulate the bone-TME, we excluded patients with neuroendocrine features as these patients develop osteolytic bone metastasis and represent a distinct subtype of PCa. Similar hierarchical clustering analyses were also performed in lymph node metastasis from the aforementioned mCRPC RNA-seq dataset(15) and treatment-naïve primary prostate cancers from the TCGA data set (16). There was no significant difference in survival between the M2-high versus M2-low populations in patients with lymph node metastasis (logRank P=0.7749) (**Fig. 1C**) or in patients with primary prostate cancer (logRank P=0.6810) (**Fig. 1C**). These observations suggest that M2-like macrophages play a more critical role in PCa bone metastasis than in lymph node metastasis or primary cancer.

**Figure 1.**
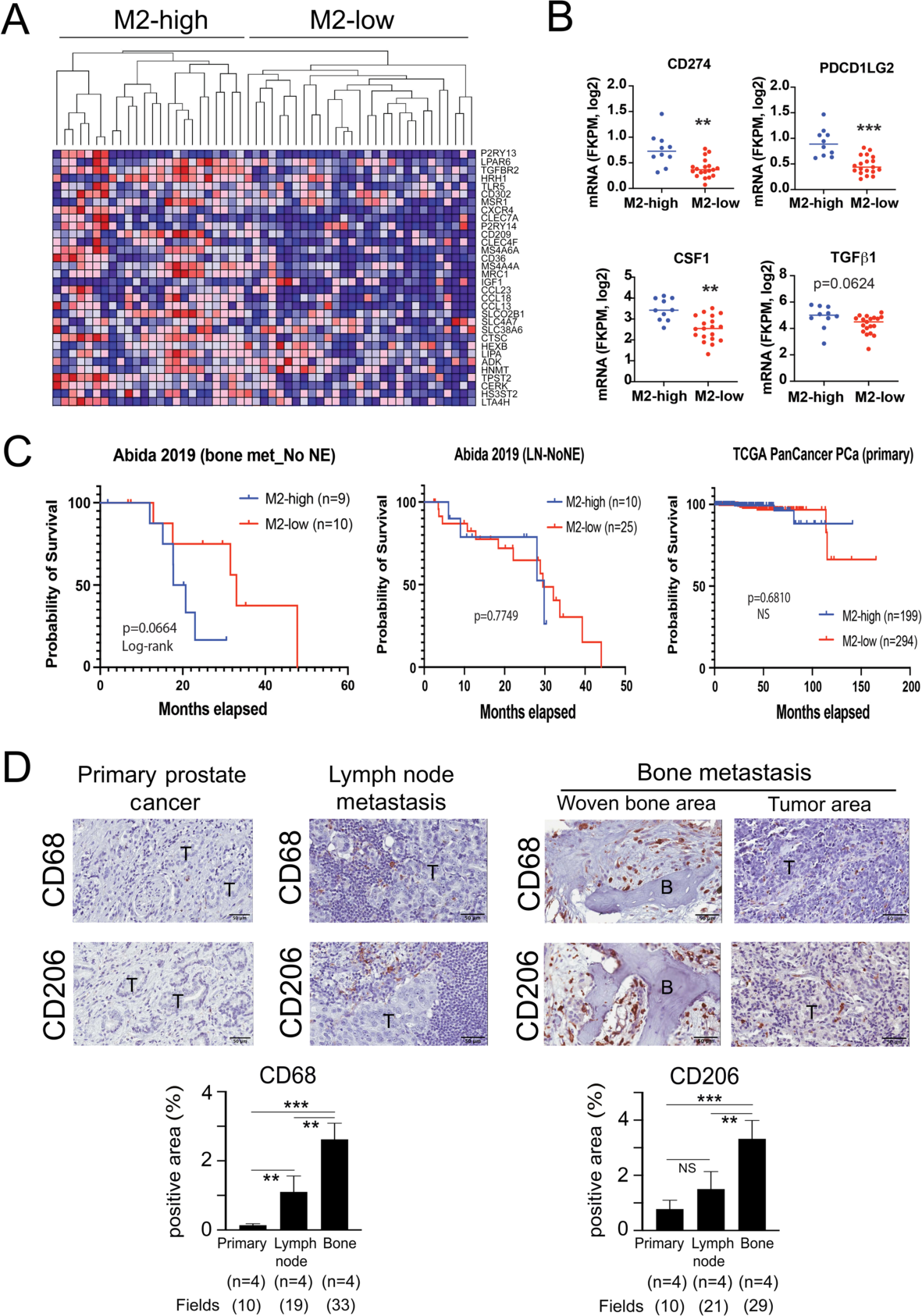
High levels M2-like macrophage in bone metastatic prostate cancer. (A) Analysis of RNA-seq dataset of bone metastatic prostate cancer using M2-like macrophage gene signature. Samples were classified into M2-high and M2-low using hierarchical clustering. (B) Higher expression of several Immunosuppressive genes in M2-high bone metastasis samples compared to M2-low samples. (C) Patients with M2-high signature in bone metastasis group, but not in lymph node metastasis or primary prostate cancer groups, had shorter survival. Neuroendocrine subtype tumors were excluded from the analysis. (D) Immunohistochemistry of CD68^+^ and CD206^+^ cells in specimens from bone metastasis (n=4), lymph node metastasis (n=4), or primary prostate cancer (n=4). CD68 or CD206 staining was quantified using ImageJ. Images from each sample were taken and quantified using ImageJ. Fields, number of regions under microscope (x200) selected for quantification. **p<0.01, ***p<0.001

### CD206^+^ macrophages are enriched in human prostate cancer bone biopsy specimens

We next performed IHC for CD68, a general macrophage marker, and CD206, a marker for M2-like macrophages, on human bone metastasis, lymph node metastasis, and primary prostate cancers. We found that bone metastasis specimens, including areas of woven bone and tumor, had higher levels of CD68^+^ macrophage infiltration (2.62 ± 0.47%) compared to lymph node metastasis (1.10 ± 0.46 %, p=0.006) or primary prostate cancers (0.14 ± 0.05 %, p<0.001) as shown by quantification of CD68^+^ cells using ImageJ (**Fig. 1D, Supplementary Fig. 1**). There was also a significant difference in frequency of CD68^+^ cells between lymph node metastasis and primary PCa (p=0.004). IHC of the specimens with anti-CD206 antibody also showed a higher level of CD206^+^ macrophage infiltration in bone metastasis (3.32 ± 0.67%) compared to lymph node metastasis (1.50±0.63%, p=0.008) or primary prostate cancers (0.78± 0.32, p<0.001). There was no significant difference in the frequency of CD206^+^ macrophages between lymph node metastasis and primary PCa (p=0.09). These results suggest M2-like macrophages are uniquely enriched in the bone-TME.

### Tumor-induced bone increases F4/80^+^ and CD206^+^ macrophage infiltration

To examine whether tumor-induced bone contributes to the increased density of CD206^+^ cells in the bone-TME, we performed IHC staining on MDA PCa-118b xenograft, an osteogenic PDX derived from a bone biopsy of a patient with castrate-resistant PCa (9). F4/80 was used as a general mouse macrophage marker. We found a high density of F4/80^+^ and CD206^+^ cells around the tumor-induced bone areas (**Fig. 2A**), consistent with those observed in human bone metastasis specimens. We have previously shown that MDA-PCa-118b expresses BMP4, leading to ectopic bone formation (7). To examine whether tumor-induced bone plays a role in increased M2-like macrophage infiltration seen in bone metastasis, we generated an osteogenic PCa model by ectopically expressing BMP4 in C4-2b PCa cells, a non-osteogenic cell line. As we previously reported (13), BMP4 overexpression in C4-2b cells led to woven bone formation in the subcutaneous tumors (**Fig. 2B**). When tumors were immune-stained for F4/80 or CD206, the densities of F4/80^+^ macrophages and CD206^+^ macrophages in the tumor areas of C4-2b-vec and C4-2b-BMP4 were similar (**Fig. 2B**). However, levels of macrophage infiltration (F4/80^+^ macrophage or CD206^+^ macrophages) were significantly higher in the areas adjacent to tumor-induced bone of C4-2b-BMP4 tumors (**Fig.2B**). We also generated osteogenic MycCaP-BMP4 tumors by transfecting mouse BMP4 cDNA into MycCaP cells. Higher levels of F4/80^+^ and CD206^+^ macrophage infiltration was observed in the MycCaP tumors compared to C4-2b tumors as shown by IHC staining (**Fig. 2C**), suggesting intrinsic variations in the levels of macrophage infiltration in tumors with different genetic backgrounds. Similar to C4-2b-BMP4 tumors, MycCaP-BMP4 tumors also evidenced higher levels of F4/80^+^ and CD206^+^ staining macrophages compared to MycCaP tumors (**Fig. 2C**). However, there was no significant difference in macrophage infiltration between the tumor-induced bone and tumor areas in MycCaP-BMP4 tumors (**Fig. 2C**), possibly due to high levels of macrophage infiltration in this genetic background.

**Figure 2.**
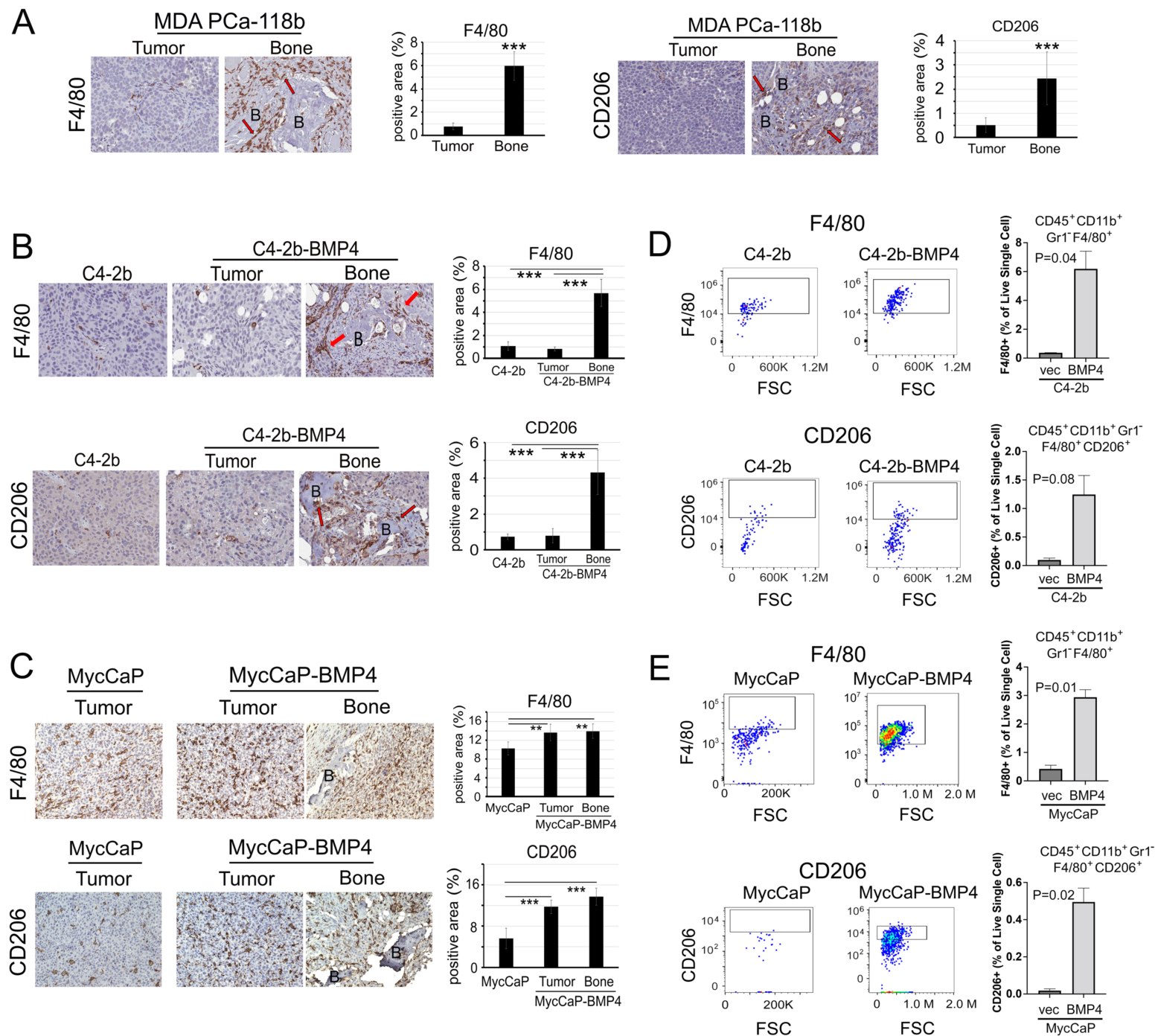
Osteogenic tumors have higher tumor-associated macrophages. (A) IHC of F4/80 (left panel) or CD206 (right panel) in MDA-PCa-118b tumors. Macrophages were enriched in areas with tumor-induced bone. (B) IHC of F4/80 or CD206 in C4-2b-vec and C4-2b-BMP4 tumors. C4-2b-BMP4 tumors had higher levels of macrophages compared to C4-2b-vec tumor and were enriched in areas with tumor-induced bone. (C) IHC of F4/80 or CD206 in MycCaP and MycCaP-BMP4 tumors. MycCaP-BMP4 tumors had higher levels of macrophages compared to MycCaP in both tumor and bone-containing areas. (D) FACS analysis of macrophages in C4-2b-vec and C4-2b-BMP4 tumors. Equal weights of tumors were digested and the cells were subjected to FACS. The levels of CD45^+^CD11b^+^Gr1^−^F4/80^+^ and CD45^+^CD11b^+^Gr1^−^F4/80^+^CD206^+^ cells in C4-2b-BMP4 tumors were higher compared to those in C4-2b-vec tumors. Average of 2 biological replicates from one experiment is shown. The study was repeated 3 times with similar results. (E) FACS analysis of macrophages in MycCaP and MycCaP-BMP4 tumors as in (D). Average of 2 biological replicates from one experiment is shown. The study was repeated 3 time with similar results. B, tumor-induced bone. Arrow, F4/80^+^ or CD206^+^ macrophages.

To further confirm the IHC results, we performed fluorescence activated cell sorting (FACS) to characterize myeloid subsets using CD45^+^ (pan-immune marker), CD11^+^ (pan-myeloid marker), GR1^−^ (granulocyte marker), F4/80^+^ (general mouse macrophage marker) and CD206^+^ (M2-like macrophage marker) in C4-2b and C4-2b-BMP4 tumors. Equal weights of tumors generated from C4-2b-vec control and C4-2b-BMP4 were digested, labeled with antibodies, and subjected to FACS analysis. We found that the levels of CD45^+^CD11b^+^Gr1^−^F4/80^+^ and CD45^+^CD11b^+^Gr1^−^F4/80^+^CD206^+^ cells were higher in C4-2b-BMP4 tumors than in control C4-2b-vec tumors (**Fig. 2D**). The p value for the CD206^+^ population in the total C4-2b and C4-2b-BMP4 tumors was 0.08, due to very few CD206^+^ cells in the C4-2b-Ctrl tumors that resulted in high variations in cell counts after the sample preparation for FACS analysis. In MycCaP-BMP4 tumors, we also observed a significantly higher number of CD45^+^CD11b^+^Gr1^−^F4/80^+^ and CD45^+^CD11b^+^Gr1^−^F4/80^+^CD206^+^ cells compared with those in MycCaP tumors (**Fig. 2E**). Taken together, these results suggest that osteogenic tumors have a higher number of tumor-associated M2-like macrophages.

### Conditioned medium from EC-OSB hybrid cells enhances the recruitment of M2-like macrophages

We have previously shown that PCa-induced bone is produced through endothelial-to-osteoblast (EC-to-OSB) transition by PCa-secreted BMP4 (13). In this process, BMP4 induces endothelial cells to first transition into EC-OSB hybrid cells that then undergo osteoblast differentiation leading to mineralization (17). To examine the role of EC-OSB hybrid cells in macrophage recruitment, 2H11 endothelial cells were treated with 100 ng/mL BMP4 for 48 hrs to generate EC-OSB hybrid cells (**Fig. 3A**). Conditioned medium (CM) from 2H11 or EC-OSB cells were assessed for their chemotactic effect on immortalized bone marrow-derived macrophages (iBMDM), RAW264.7 cells, and mouse bone-marrow-derived macrophages (BMDM). iBMDM and RAW264.7 cells were polarized to M2-like macrophages by incubating with IL-4 for 24 hr as shown by the upregulation of M2 macrophage markers *arginase1 (Arg1), Mrc1(CD206)* and *Clec10a*, and to M1 macrophages by incubating with lipopolysaccharide (LPS) as shown by the upregulation of M1 macrophage markers *Tnf, Nos2,* and *Ccl2* (**Supplementary Fig. 2**). Similarly, BMDM were polarized to M2-like macrophage by incubating with IL-4 for 72 h as shown by the upregulation of M2 macrophage markers (**Supplementary Fig. 2**). Boyden Chamber migration assay was used to examine the effect of CM on macrophage recruitment. EC-OSB CM, but not control 2H11 CM or BMP4, which was used to induce EC-to-OSB transition, dramatically promoted the migration of M2-polarized, but not M1-polarized iBMDM (**Fig. 3B**). Similar results were observed with RAW264.7 macrophages (**Fig. 3C**). These effects were further confirmed using BMDMs that were polarized to M2 phenotype by IL-4 (**Fig. 3D**). These results suggest that factors secreted from EC-OSB cells recruit M2-like macrophages.

**Figure 3.**
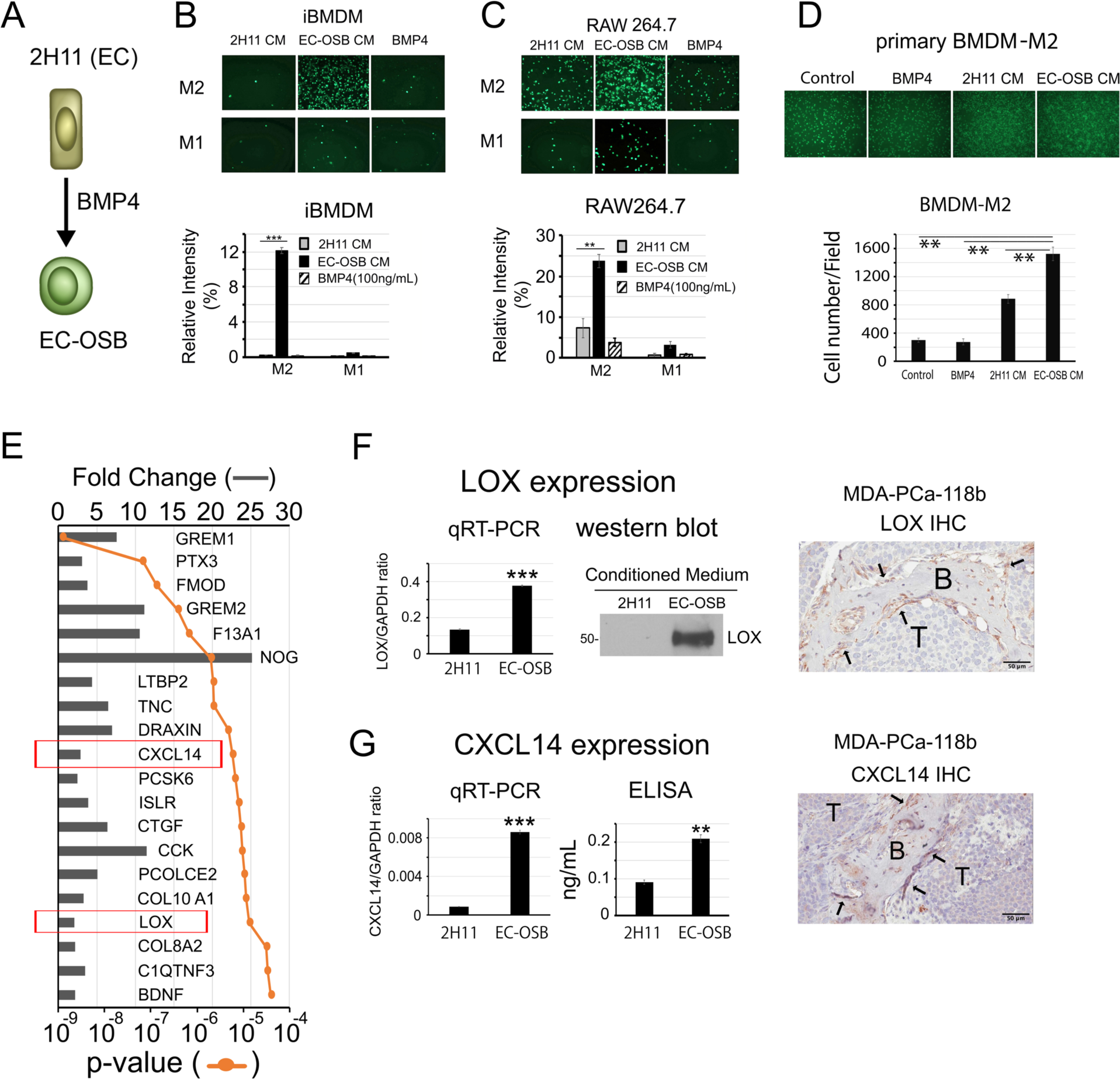
Conditioned medium from EC-OSB hybrid cells enhances the recruitment of M2-like macrophages. (A) EC-OSB hybrid cells were generated by treating 2H11 endothelial cells with 100 ng/mL BMP4 for 48 hrs. The conditioned media from 2H11 cells or EC-OSB hybrid cells was collected for migration assay. (B) Boyden Chamber migration assay for M1 or M2-polarized iBMDM, (C) M1 or M2-polarized RAW264.7, and (D) M2-polarized BMDM macrophages. EC-OSB CM promoted the migration of M2-polarized, but not M1-polarized macrophages. (E) RNAseq analysis of EC-OSB cell-secreted factors from 2H11 cells treated with or without BMP4. The top 20 genes with p< 10^−4^ were shown. The values were expressed as the fold change between EC-OSB hybrid cells and 2H11 cells. (F) Confirmation of LOX and (G) CXCL14 expression in EC-OSB cells by qRT-PCR, Western blot or ELISA, and IHC of MDA PCa 118b tumor. B, bone; T, tumor. ** p<0.01; *** p<0.001.

### LOX and CXCL14 in EC-OSB CM are candidate factors for M2-macrophage recruitment

To identify secreted factors in EC-OSB CM that led to M2-macrophage recruitment, RNAseq analysis of RNA from 2H11 and EC-OSB cells (GSE168321) was performed and analyzed for differentially expressed secreted factors (**Fig. 3E**). The most abundantly secreted factors, including gremlin 1, gremlin 2, and Noggin, are BMP inhibitors, reflecting a negative feedback mechanism following BMP4 treatment. Among other secreted factors upregulated in EC-OSB CM, lysyl oxidase (LOX) and CXCL14 have previously been shown to recruit macrophages (18,19). LOX was shown to enhance macrophage migration in glioblastoma (18), and CXCL14 was shown to recruit M2-like macrophages in brown fat (19). Upregulation of LOX and CXCL14 in the EC-OSB cells was confirmed by qRT-PCR (**Fig. 3F, G**). Western blot and ELISA further showed that both LOX and CXCL14 proteins were secreted into the EC-OSB CM (**Fig. 3F, G**). Furthermore, IHC of MDA-PCa-118b tumors with anti-LOX or anti-CXCL14 localized the expression of LOX and CXCL14 in EC-OSB hybrid cells rimming the tumor-induced bone in MDA-PCa-118b tumor (**Fig. 3F, G, Arrows**). These findings suggest that LOX and CXCL14 secreted from EC-OSB cells during tumor-induced bone formation are involved in M2-like macrophage recruitment.

### LOX and CXCL14 secreted by EC-OSB hybrid cells promote macrophage migration

To examine the function of LOX as macrophage chemoattractant in vitro, medium supplemented with recombinant LOX protein (200 ng/mL) was used in a transwell assay. LOX increased M2-polarized, but not M1-polarized, iBMDM, RAW264.7 and BMDM macrophage migration compared to control medium (**Fig. 4A**). Knockdown of *Lox* using shRNA resulted in a significant decrease in the *Lox* mRNA expression by qRT-PCR and protein in EC-OSB cell lysates by western blot (**Fig. 4B**). Using conditioned media (CM) from BMP4-treated shLox or control shvec-transfected 2H11 cells for transwell assay, knockdown of Lox led to a decrease in the migration of M2-polarized iBMDM and RAW264.7 cells (**Fig. 4C**). Knockdown of *Lox* did not have significant effects on BMP4-mediated EC-to-OSB transition as measured by qRT-PCR analysis of osteocalcin (*Bglap*) mRNA expression (**Supplementary Fig. 3A**) and on mineralization as shown by Alizarin red staining (**Supplementary Fig. 3B**), suggesting that Lox is not essential for EC-to-OSB transition. A similar approach was used to examine the function of CXCL14 as a macrophage chemoattractant *in vitro*. Recombinant CXCL14 protein (100 ng/mL) increased migration of M2-polarized macrophages compared to control medium (**Fig. 4D**) and shRNA knockdown of *Cxcl14* in 2H11 cells led to a decrease in the migration of M2-polarized iBMDM and RAW264.7 cells (**Fig. 4E, F**). Knockdown of *Cxcl14* did not have significant effects on BMP4-mediated EC-to-OSB transition (**Supplementary Fig. 3A, B**). Together, these results suggest that LOX and CXCL14 secreted during EC-to-OSB transition play a role in recruiting M2-like macrophages into the bone-TME (**Fig. 4G**).

**Figure 4.**
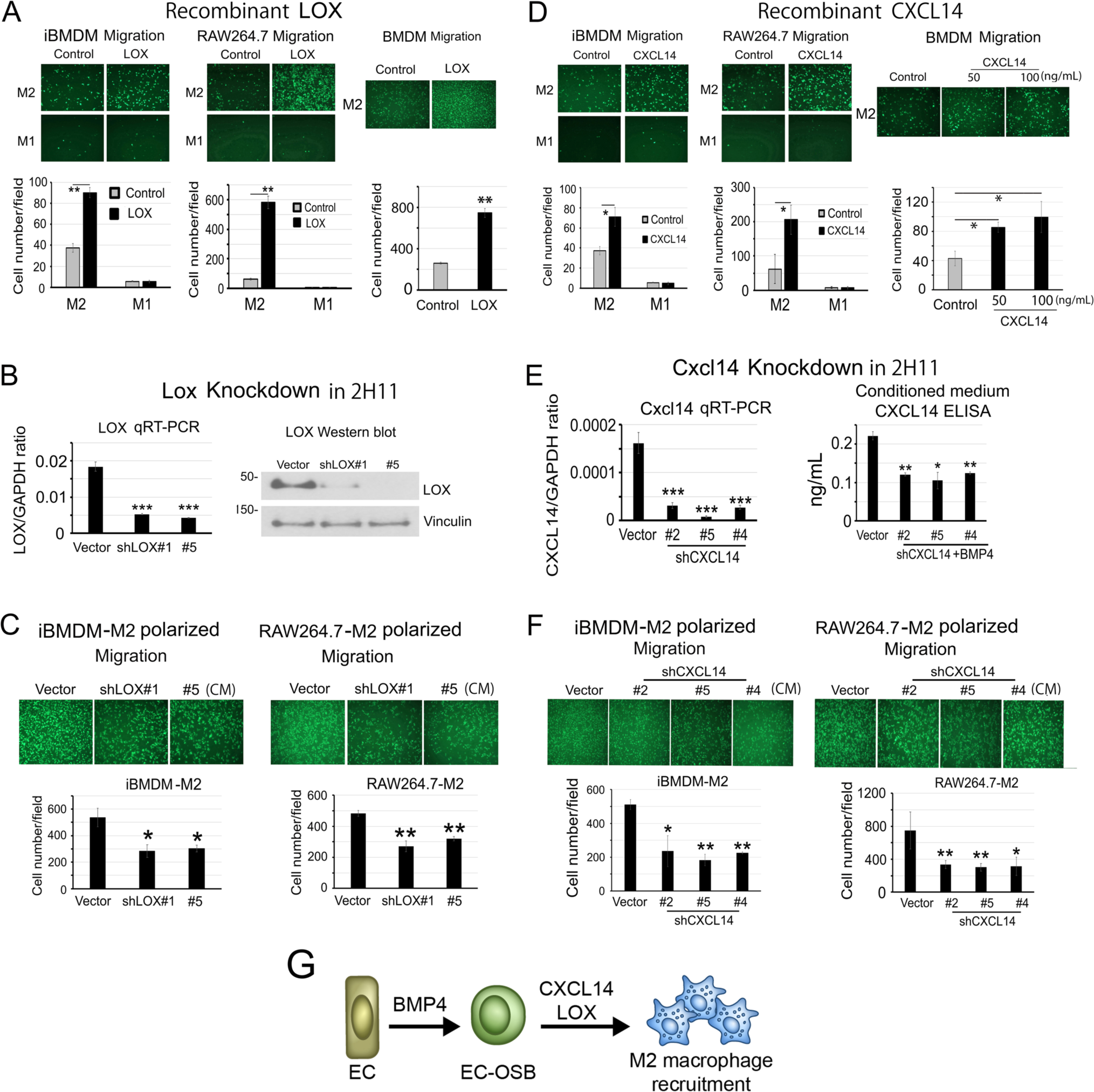
CXCL14 and LOX proteins secreted from EC-OSB hybrid cells promote macrophage migration. (A) Recombinant LOX protein (200 ng/mL) increased the migration of M2-polarized, but not M1-polarized, iBMDM, RAW264.7 and BMDM macrophages compared to medium control medium. n = 3– 4 biological replicates. (B) Knockdown of *Lox* decreased the mRNA in EC-OSB cells and protein in EC-OSB conditioned medium. (C) Conditioned media from EC-OSB cells expressing *Lox* shRNA (sh*Lox*) reduced the migration of M2-polarized iBMDM or RAW264.7 cells compared with shvector EC-OSB CM. (D) Recombinant CXCL14 protein (100 ng/mL) increased the migration of M2-polarized, but not M1-polarized, iBMDM, RAW264.7 and BMDM macrophages compared to control medium. n = 3–4 biological replicates. (E) Knockdown of *CXCL14* decreased the mRNA in EC-OSB cells and protein in EC-OSB conditioned medium. (F) Conditioned media from EC-OSB cells expressing *CXCL14* shRNA (sh*Cxcl14*) reduced the migration of M2-polarized iBMDM or RAW264.7 cells compared with shvector EC-OSB CM. (G) LOX and CXCL14 secreted from EC-OSB cells plays a role in recruiting M2-macrophages. Error bars indicate SD. *p < 0.05, **p < 0.01, **p < 0.001, Student’s t test.

### Inhibition of EC-to-OSB transition by palovarotene or LDN193189 decreases CD206^+^ TAM infiltration

We next examined whether inhibiting PCa-induced bone formation would reduce CD206^+^ macrophage infiltration in tumors. We first examined the effect of pharmacological inhibitiors of EC-to-OSB transition on the recruitment of M2-like macrophages into the bone-TME. Our previous studies showed that treatment of mice with MDA-PCa-118b tumors with LDN193189, a BMP receptor kinase inhibitor, reduced tumor-induced bone formation (7). Recently, we also showed that the RARγ agonists, including palovarotene and all-trans-retinoic acid, inhibited BMP4-mediated EC-to-OSB transition through promoting pSmad1 degradation (20) (**Fig. 5A**). To examine whether inhibition of tumor-induced EC-to-OSB transition could reduce M2-like macrophage infiltration, mice with MDA-PCa-118b tumors were treated with LDN193189 or all-trans retinoic acid (ATRA). IHC analysis showed that LDN194189 or ATRA treatment significantly reduced CD206^+^ macrophages in MDA-PCa-118b tumors (**Fig. 5B**). Similarly, treatment of C4-2b-BMP4 tumors with LDN193189 or palovarotene also significantly reduced CD206^+^ macrophages (**Fig. 5C, Supplementary Fig. 4**). Of note, palovarotene and LDN193189 had minimal effects on macrophage proliferation *in vitro* as measured by MTS assay (**Supplementary Fig. 5**). These results suggest that inhibition of tumor-induced bone formation decreases CD206^+^ macrophage infiltration in osteogenic tumors.

**Figure 5.**
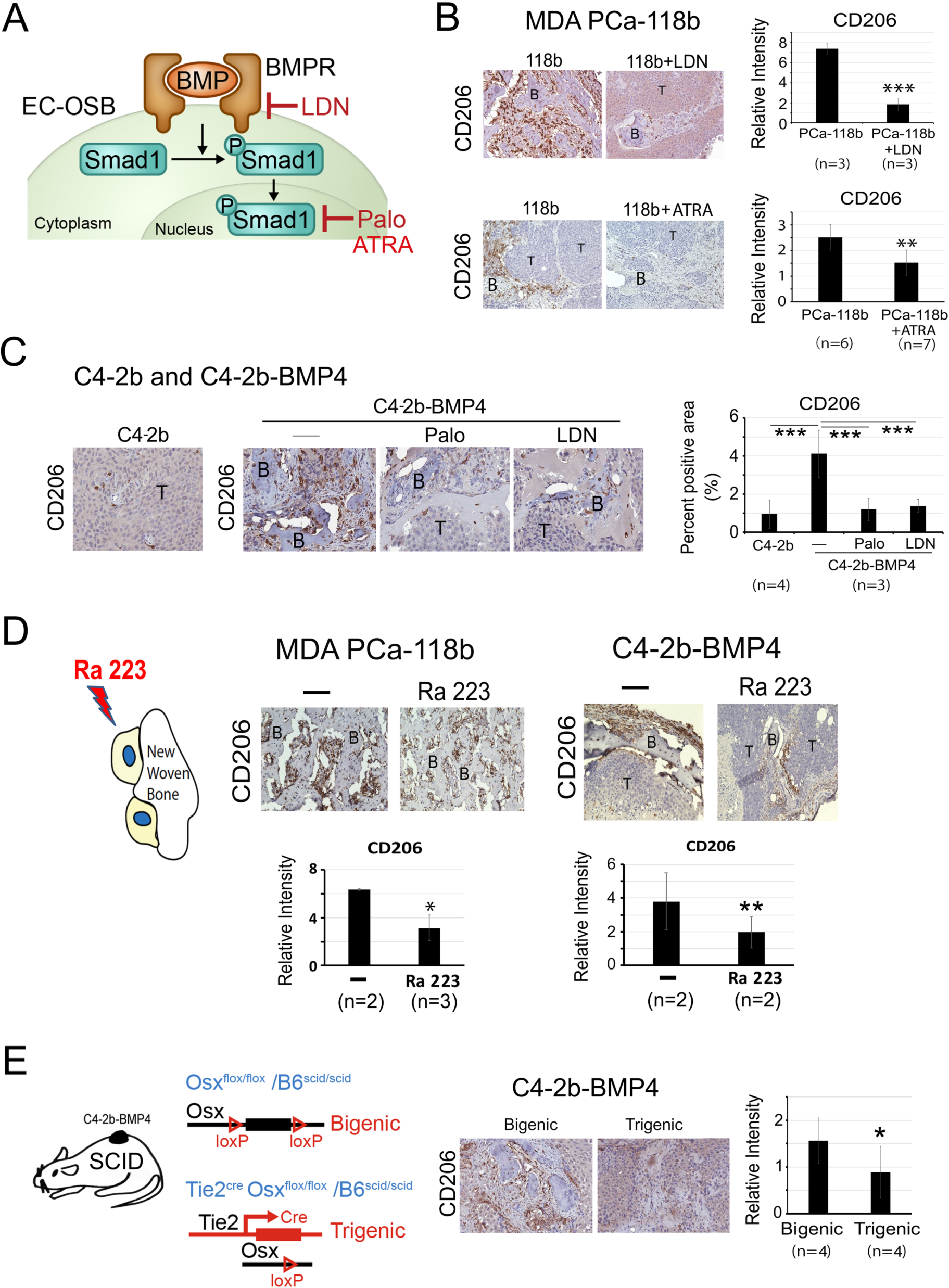
Inhibition of tumor-induced bone formation in C4-2b-BMP4 or MDA PCa-118b tumors reduces CD206^+^ tumor-associated macrophages. (A) LDN193189, palovarotene (Palo), and ATRA inhibit components of the BMP4 signal pathway. (B) Inhibition of tumor-induced bone by LDN193189 (3 mg/kg/twice a day, intraperitoneally) or ATRA (10 mg/kg/day, orally) in MDA PCa-118b tumors reduces CD206^+^ macrophage. (C) IHC of CD206 in C4-2b-vec or C4-2b-BMP4 tumors from mice treated with or without palovarotene (Palo)(2 mg/kg/day, orally) or LDN193189 (3 mg/kg/day, intraperitoneally). (D) IHC of CD206 in MDA PCa-118b and C4-2b-BMP4 tumors treated with or without Ra223. (E) CD206^+^ macrophages in C4-2b-BMP4 tumors from trigenic mice are lower compared to those from bigenic mice. Levels of CD206^+^ macrophage were quantified by image J. B, tumor-induced bone. T, tumor. N denotes the number of tumors examined. Multiple random fields were selected for quantification using Image J.

### Radium-223 treatment decreases F4/80^+^ and CD206^+^ TAM infiltration to MDA-PCa-118b or C4-2b-BMP4 tumors in SCID mice

Radium-223 (Ra223) is a high energy α-emitting calcium mimetic that preferentially localizes to hydroxyapatite in newly formed matrix within osteogenic metastases and induces double strand DNA breaks in adjacent cells (21). Ra223 preferentially targets newly formed bone and is an approved agent in the treatment of prostate cancer patients with bone metastasis based on an improvement in overall survival (22). We next examined whether Ra223 reduced M2-like TAMs in osteogenic tumors. MDA PCa-118b cells were injected subcutaneously into SCID mice. Tumor bearing mice were then treated with or without one dose (300 kBq/kg) of Ra223. Immunohistochemical analysis of MDA-PCa-118b tumors showed that Ra223 treatment reduced CD206^+^ TAMs in MDA PCa-118b tumors (**Fig. 5D**). Similar results were observed in osteogenic C4-2b-BMP4 tumors (**Fig. 5D**). Together, these results suggest that targeting EC-OSB cells with Ra223 in the bone-forming tumors MDA-PCa-118b or C4-2b-BMP4 decreases levels of CD206^+^ macrophages within the bone-TME.

### Reduction of tumor-induced bone in tumors implanted in an endothelial-specific osterix deletion model (*Tie2^cre^;Osx^f/f^;Scid^+/+^*) decreases CD206^+^ macrophage infiltration

We next examined whether reducing PCa-induced bone formation had an effect on CD206^+^ macrophage infiltration in osteogenic tumors generated in a genetically modified mouse model. In our previous studies, we generated a genetically modified mouse model (*Tie2*^cre^/*Osx*^f/f^) in a SCID background with endothelial-specific deletion of *osterix* (*OSX*), a cell fate determinant gene that controls the development of the osteoblast lineage (23). We used this model to examine the role of EC-to-OSB transition in PCa-induced osteogenesis (**Fig. 5E**). The tumor-induced bone in C4-2b-BMP4 tumors was significantly reduced when tumors were implanted in (<u>*Tie2-Cre;Osx^f/f^;Scid^+/+^*</u>; **<u>trigenic</u>**) mice, in which OSX in endothelial cells was specifically deleted, compared to those implanted in control (<u>*Osx^f/f^;Scid^+/+^*</u>; **bigenic**) mice (13). We examined whether a reduction of tumor-induced bone formation in the trigenic mice had an effect on tumor-associated macrophages. IHC analysis of C4-2b-BMP4 tumors from trigenic mice showed a significant decrease of CD206^+^ macrophages compared with those from control bigenic mice (**Fig. 5E**). These results suggest that reducing tumor-induced bone formation through genetic deletion of *Osx* in endothelial cells reduces CD206^+^ macrophage infiltration in the bone-TME.

### RNAseq analysis of MycCaP and MycCaP-BMP4 bulk tumors in FVB mice revealed enrichment of an M2-like macrophage signature in osteogenic tumors

To define the molecular differences between MycCaP and MycCaP-BMP4 tumors, we performed bulk RNAseq on total RNA isolated from MycCaP or MycCaP-BMP4 tumors (GSE241343). MycCaP or MycCaP-BMP4 cells were implanted in immune-competent FVB mice, the mouse strain in which MycCaP tumors were originally generated (24,25). Because MycCaP-BMP4 cells that possessed a selection marker (e.g., GFP) could not grow in FVB mice, likely due to an immune response against the selection marker, the marker negative MycCaP-BMP4 cells for these experiments were generated by dilutional cloning. MycCaP-BMP4 tumors in FVB mice showed tumor-induced bone formation and recruitment of F4/80^+^ and CD206^+^ macrophages to the tumor-induced bone area (**Supplementary Fig. 6**), similar to those observed in SCID mice.

Using the M2-gene signature shown in Fig. 1A, GSEA analysis of GSE241343 RNAseq data demonstrated an enrichment of an M2-like macrophage signature in MycCaP-BMP4 tumors compared to those from MycCaP tumors (**Fig. 6A**). Next, using CIBERSORT deconvolution algorithms (26) to analyze for M0, M1, or M2 populations, M2-like macrophages were enriched in MycCaP-BMP4 compared to MycCaP tumors (**Fig. 6B**), while M0 and M1 macrophages were enriched in MycCaP tumors compared to MycCaP-BMP4 tumors (**Fig. 6B**). Consistent with the CIBERSORT results, DEG analysis comparing MycCaP-BMP4 tumors to MycCaP tumors showed that M2 markers *Mrc1*, *Clec10a*, *Il10*, *Cxcl13* (27), *Il1r2* (28), *Vegfa*, and *Vegfd* were upregulated while M1 markers *Tnf*, *Il16* (29), and *Ifnlr1* (30) were downregulated (**Supplementary Fig. 7A**). Furthermore, qRT-PCR confirmed upregulation of M2-macrophage markers, including *Mrc1* (*CD206*), *IL10* and *Clec10a*, in MycCaP-BMP4 tumor (**Supplementary Fig. 7B**), while M1-macrophage markers *Ccr2*, *Nos2*, *Ccl2*, and *Il1β*, were highly expressed in MycCaP tumors (**Supplementary Fig. 7B**).

**Figure 6.**
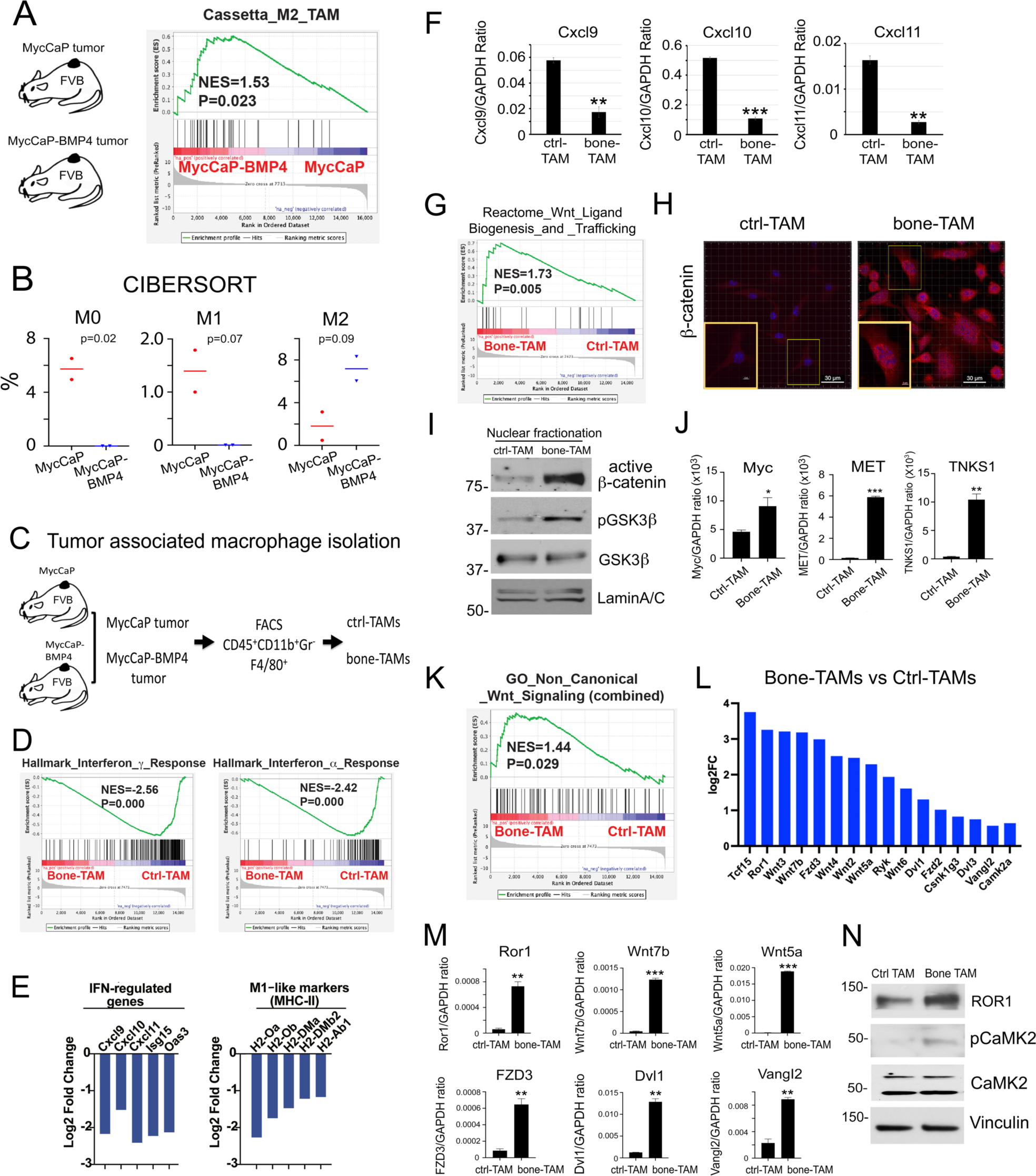
RNAseq analysis of tumor-associated macrophages revealed upregulation of M2 macrophage-related and downregulation of M1 macrophage-related genes in MycCaP-BMP4 tumors. (A) RNAseq analysis of MycCaP and MycCaP-BMP4 tumors implanted in FVB mice. An M2-like macrophage signature was enriched in MycCaP-BMP4 tumors compared to those in MycCaP tumors. (B) Analysis of RNAseq data from MycCaP vs MycCaP-BMP4 tumors using CIBERSORT. MycCaP tumors have higher levels of M0 and M1 macrophage signatures, while MycCaP-BMP4 tumors have higher levels of an M2-like macrophage signature compared to MycCaP tumors. (C) Schema for isolating tumor associated macrophages (TAMs) from MycCaP and MycCaP-BMP4 tumors. (D) GSEA analysis showed interferon gamma and alpha signaling were suppressed in bone-TAMs. (E) Decreases in the expression levels of IFN-regulated genes and MHC-II genes in bone-TAMs compared to ctrl-TAMs. (F) qRT-PCR of RNAs isolated from ctrl-TAMs and bone-TAMs. Several M1 macrophage markers, including Cxcl9, Cxcl10, Cxcl11, were downregulated in bone-TAMs. (G) GSEA analysis showed Wnt signaling was activated in bone-TAMs compared to ctrl-TAMs. (H) Immunofluorescence using antibody against active β-catenin showed an increase of β-catenin in the nucleus of bone-TAM compared to ctrl-TAM. Inset, β-catenin in the nucleus of a bone-TAM. (I) Western blot of the nuclear fractions of ctrl-TAMs and bone-TAMs. The levels of active β-catenin and pGSK3β were higher in bone-TAM compared with ctrl-TAMs. (J) qRT-PCR showed that the mRNA levels of several downstream canonical Wnt pathway target genes were upregulated, but some were not, in bone-TAMs compared to ctrl-TAMs. (K) GSEA analysis for genes involved in non-canonical Wnt signaling showed upregulation of non-canonical Wnt pathways in bone-TAMs compared to ctrl-TAMs. (L) Fold changes of genes in non-canonical Wnt signaling. (M) qRT-PCR confirmed the upregulation of several genes in non-canonical Wnt signaling. (N) Western blot confirmed the increased levels of ROR1 and its downstream pCAMKII proteins in bone-TAMs compared to ctrl-TAMs.

### RNAseq analysis of tumor-associated macrophages (TAMs) revealed a reduction of an IFN-TAM gene signature and activation of canonical and non-canonical Wnt pathways in MycCaP-BMP4 tumors

To characterize the key molecular features of macrophages recruited to the TME of MycCaP and MycCaP-BMP4 tumors, CD45^+^CD11b^+^Gr1^−^F4/80^+^ macrophages were isolated by FACS from MycCaP/FVB tumors (ctrl-TAMs) and MycCaP-BMP4/FVB subcutaneous tumors (bone-TAMs) (**Fig. 6C**) and RNAseq analysis was performed (GSE 241344). GSEA showed suppression of both IFN gamma and alpha pathways in bone-TAMs compared to ctrl-TAMs (**Fig. 6D**). Consistent with the GSEA findings, IFN-regulated genes including *Cxcl9*, *Cxcl10*, *Cxcl11*, *Isg15*, and *Oas3*, were downregulated in bone-TAMs compared to ctrl-TAMs (**Fig. 6E**). In addition, several MHC-II genes (e.g., *H2-Oa*, *H2-Ob*, and *H2-DMa*) were significantly reduced in bone-TAMs compared to ctrl-TAMs (**Fig. 6E**). Recent studies using single cell RNAseq have revealed significant heterogeneity in TAMs and a proposed classification into functionally distinct subsets has been proposed, including interferon-primed TAMs (IFN-TAMs), immune regulatory TAMs (Reg-TAMs), inflammatory cytokine-enriched TAMs (Inflam-TAMs), lipid-associated TAMs (LA-TAMs), pro-angiogenic TAMs (Angio-TAMs), RTM-like TAMs (RTM-TAMs), and proliferating TAMs (Prolif-TAMs) (31). Using these TAM subset gene signatures, we found that IFN-TAM gene signature was dramatically downregulated in bone-TAMs (**Supplementary Fig. 7C**). Importantly, downregulation of IFN-TAM-associated genes (*Cxcl9*, *Cxcl10*, *Cxcl11*, *H2-Oa*, and *H2-Ob*) was confirmed by qRT-PCR in isolated bone-TAMs compared to ctrl-TAMs (**Fig. 6F, Supplementary Fig. 7D**). These results suggest that osteogenic tumors have a reduction of the IFN-TAMs in the bone-TME compared to non-osteogenic tumors.

GSEA showed that pathways related to WNT signaling were among the top 10 upregulated pathways in bone-TAMs (KEGG, Reactome, WikiPathways) (**Fig. 6G, Supplementary Fig. 7E & Supplementary Table 1**). In lung cancer, Sarode et al. (32) showed that macrophage-specific β-catenin-mediated Wnt signaling is centrally involved in the phenotypic transition of M1-like to M2-like TAMs. To consider this possibility in prostate cancer, we next examined whether the Wnt pathway is activated in bone-TAMs using our model. Immunofluorescence showed an increase of β-catenin, the key signal transducer of Wnt pathway, in both the nucleus and cytoplasm of FACS sorted bone-TAMs compared to ctrl-TAMs (**Fig. 6H**). Furthermore, Western blot showed increased nuclear β-catenin and pGSK3β, which regulates the phosphorylation and stability of β-catenin, in bone-TAMs compared with ctrl-TAMs (**Fig. 6I**). qRT-PCR showed that the mRNA levels of the downstream canonical Wnt pathway target genes, *Myc*, *MET*, tankyrase (*TNKS1*), but not *TNKS2* and *CCND1*, were upregulated in bone-TAMs compared to ctrl-TAMs (**Fig. 6J**, **Supplementary Fig. 7F**). Besides the canonical Wnt pathway, GSEA analysis also showed upregulation of the non-canonical Wnt pathway in bone-TAMs compared to ctrl-TAMs (**Fig. 6K**). Genes in non-canonical Wnt pathway that were upregulated in bone-TAMs included *Ror1*, *Wnt7b*, *Wnt5a*, *FZD3*, *Dvl1*, and *Vangl2* (33–35) (**Fig. 6L**). Of note, several Wnt ligands including *Wnt2, 3, 4, 5a, 6*, and *7b*, were also upregulated (**Fig. 6L**), suggesting autocrine activation of the Wnt pathways in bone-TAMs. qRT-PCR confirmed upregulation of genes involved in non-canonical Wnt signaling in bone-TAMs compared to ctrl-TAMs (**Fig. 6M**). Western blot confirmed increased levels of ROR1 and its downstream target pCAMKII, in bone-TAMs compared to ctrl-TAMs (**Fig. 6N**). These results suggest that upregulation of both canonical and non-canonical Wnt pathways represents a key molecular feature of bone-TAMs.

### EC-OSB-mediated Wnt pathway activation promotes M2 polarization

The polarization of TAMs is directly controlled by cytokines within the tumor microenvironment (36). Because bone-TAMs are enriched in osteogenic tumors, we examined whether EC-OSB hybrid cells play a role in the M2-like polarization of bone-TAMs. Mouse BMDMs were treated with EC-OSB conditioned medium (CM) for 5 days. EC-OSB CM promoted M2 polarization of BMDM as shown by increased M2 markers (**Fig. 7A**) and increased CD206 expression by Western blot (**Fig. 7B**). Similar results were obtained when RAW264.7 cells were cultured with EC-OSB conditioned medium for 48 hrs (**Supplementary Fig. 8A**).

**Figure 7.**
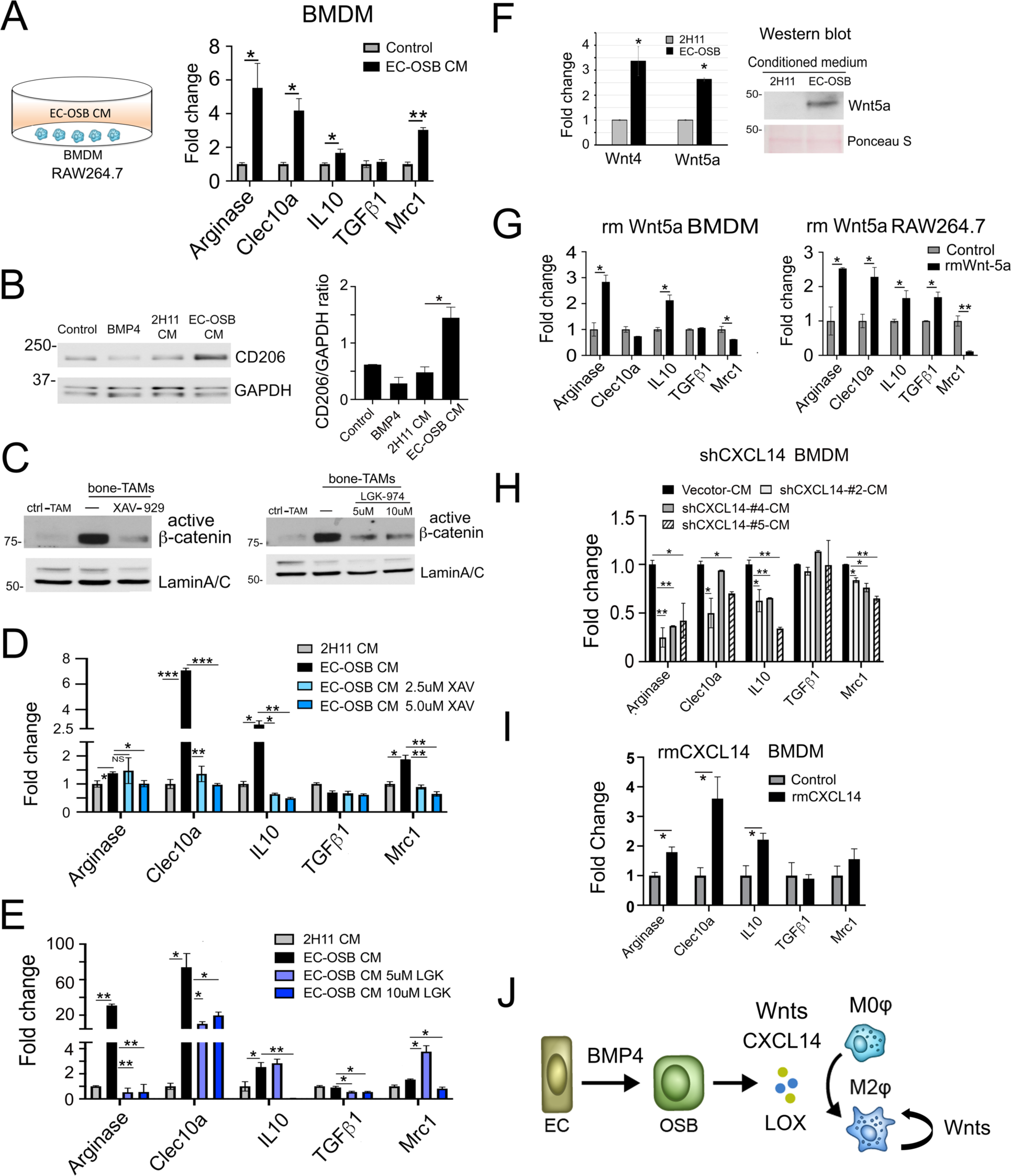
EC-OSB-secreted factors promote M2 polarization of macrophages. (A) Treatment of BMDM with EC-OSB CM upregulated markers of M2 macrophages. (B) Treatment of BMDM with EC-OSB CM led to an increase of CD206 expression. (C) Nuclear β-catenin levels in bone-TAMs were higher than those in ctrl-TAMs. Treatment of bone-TAMs with Wnt pathway inhibitor XAV-939 or LGK-974 reduced the levels of nuclear β-catenin levels in bone TAMs. (D) EC-OSB CM-mediated M2-polarization was suppressed by a Wnt pathway inhibitor XAV-939. (E) EC-OSB CM-mediated M2-polarization was suppressed by a Wnt pathway inhibitor LGK-974. (F) EC-OSB hybrid cells secrete several Wnt ligands as detected by qRT-PCR and Western blot. (G) Treatment of BMDMs or RAW264.7 cells with recombinant mouse Wnt 5a upregulated M2 marker expression. (H) Conditioned media from EC-OSB cells with knockdown of *CXCL14* (sh*Cxcl14*) reduced the M2 markers of BMDM compared with shvector EC-OSB CM. (I) Treatment of BMDMs with recombinant mouse CXCL14 protein upregulated M2 markers of BMDM. (J) Graphical summary. EC-OSB cells-secreted many factors, including Wnt ligands, CXCL14 and LOX, that promoted M2 polarization of macrophages. Bone-TAMs also express Wnts that further promoted M2 polarization in an autocrine manner. Error bars indicate SD. *p < 0.05, **p < 0.01, **p < 0.001, Student’s t test.

To examine whether the Wnt pathway was involved in EC-OSB CM-mediated M2 polarization, we used XAV-939 and LGK974, two small-molecule canonical Wnt pathway inhibitors, to block Wnt/β-catenin signaling. XAV-939 is a tankyrase inhibitor that promotes the degradation of β-catenin (37,38) and LGK974 targets Porcupine, a Wnt-specific acyltransferase (39). Treatment of bone-TAMs with XAV-939 or LGK-974 reduced levels of nuclear β-catenin levels in bone TAMs (**Fig. 7C**). BMDMs pretreated with XAV-939 before incubation with EC-OSB CM led to inhibition of the Wnt pathway as evidenced by reduced levels of M2-markers including *Clec10a*, *Il10*, and *Mrc1* (**Fig. 7D**). Similarly, pretreatment of BMDMs with LGK974 (39) also reduced levels of M2-markers (**Fig. 7E)**. These results suggest that EC-OSB-mediated Wnt pathway activation is one of the mechanisms that promote M2 polarization of TAMs in bone-TME.

### EC-OSB hybrid cells secrete WNTs, CXCL14, and LOX to promote M2 polarization

Next, we investigated factors in EC-OSB conditioned medium that activate the Wnt pathway in bone-TAMs. In our previous RNAseq analysis of 2H11 versus EC-OSB cells, we found *Wnt4* and *Wnt5a* were upregulated in EC-OSB hybrid cells compared to 2H11 cells (GSE 168321)(17). To examine the expression of Wnt ligands during EC-to-OSB transition, 2H11 endothelial cells were treated with BMP4 for 48 hrs to induce EC-to-OSB transition. qRT-PCR confirmed that *Wnt4* and *Wnt5a* were upregulated in EC-OSB cells compared to 2H11 cells, and Western blot showed upregulation of WNT5A in EC-OSB CM (**Fig. 7F**). qRT-PCR also showed that *Wnt7*, but not *Wnt6*, was upregulated in EC-OSB cells (**Supplementary Fig. 8B**), suggesting that EC-OSB cells secreted multiple Wnt ligands. Of note, WTN5A is a ligand for non-canonical Wnt pathway activation (33,35). Treatment of BMDMs or RAW264.7 with recombinant mouse WNT5A led to an increase in the expression of several M2 markers in BMDMs (*Il10* and *Arg1*) and RAW264.7 cells (e.g., *Arg1*, *Clec10a*, *Il10*, *Tgfb1*) (**Fig. 7G**), suggesting that WNT5A is involved in M2 polarization of macrophages. Interestingly, *Mrc1* expression was downregulated by WNT5A (**Fig. 7G**), suggesting that other EC-OSB secreted factors besides Wnt ligands may be also involved in M2-polarization of bone-TAMs. We found that CXCL14 secreted from EC-OSB cells played a role in the recruitment of M2-like macrophages to tumor-induced bone area (**Fig. 4D-F**). CXCL14 was also shown to function as a chemokine for M2-polarization of macrophages (19). Interestingly, Gao et al. (40) reported that CXCL14 facilitated ovarian cancer cell invasiveness through activation of Wnt/ β-catenin signaling pathway, although the mechanism is unknown. We found that knockdown of CXCL14 in EC-OSB cells reduced the EC-OSB-mediated upregulation of M2-macrophage genes (*Arg1, Clec10a, IL10, Mrc1*) in BMDMs (**Fig. 7H**) and RAW264.7 cells (**Supplementary Fig. 8C**). Furthermore, addition of recombinant mouse CXCL14 protein to BMDMs or RAW264.7 cells led to increased expression of M2-macrophage genes (**Fig. 7I, Supplementary Fig. 8D**). Similarly, knockdown of shLox in EC-OSB cells inhibited M2-polarization of BMDMs while recombinant mouse LOX protein stimulated M2-polarization of BMDMs (**Supplementary Fig. 8E-F**) and RAW264.7 cells (**Supplementary Fig. 8G-H**). These results suggest that several EC-OSB-secreted factors including WNTs, CXCL14 and LOX, play a role in EC-OSB-mediated M2-polarization (**Fig. 7J**). Furthermore, bone-TAMs also express Wnts (**Fig. 6L, M**) to further promote M2 polarization through an autocrine manner (**Fig. 7J**).

### Macrophages from C4-2b-BMP4 or MycCaP-BMP4 tumors inhibit T cell proliferation

To examine the effect of bone-TAMs on T cell function, we first examined whether bone-TAMs could suppress T cell proliferation. CD45^+^CD11b^+^Gr1^−^F4/80^+^ cells were isolated from C4-2b-vec and C4-2b-BMP4 tumors and a T-cell suppression assay was performed using CFSE-labeled mouse splenic CD8^+^ T-cells activated by anti-CD3 and anti-CD28 antibodies (CD3/CD28 Dynabeads). F4/80^+^ cells from C4-2b-BMP4 tumors significantly suppressed the proliferation of T cells compared to ctrl-TAMs from C4-2b-vec tumors (**Fig. 8A**). Next, CD45^+^CD11b^+^Gr1^−^F4/80^+^CD206^+^ cells were isolated from C4-2b-BMP4 tumors and found to inhibit T-cell proliferation (**Fig. 8B**), suggesting that reduction of T-cell proliferation is likely due to CD206^+^ M2-like TAMs. The control C4-2b-vec tumor contained too few CD45^+^CD11b^+^Gr1^−^F4/80^+^CD206^+^ cells to perform the assay, consistent with low CD206^+^ in C4-2b-vec tumors (**Fig. 2B, D**). Bone-TAMs from MycCaP-BMP4 tumors inhibited T cell proliferation compared with ctrl-TAMs (**Fig. 8C**). Similar results were observed using ctrl-TAMs and bone-TAMs isolated from MycCaP and MycCaP-BMP4 tumors generated in FVB mice (**Fig. 8D**). Thus, M2-like macrophages enriched in osteogenic tumors inhibit T-cell proliferation.

**Figure 8.**
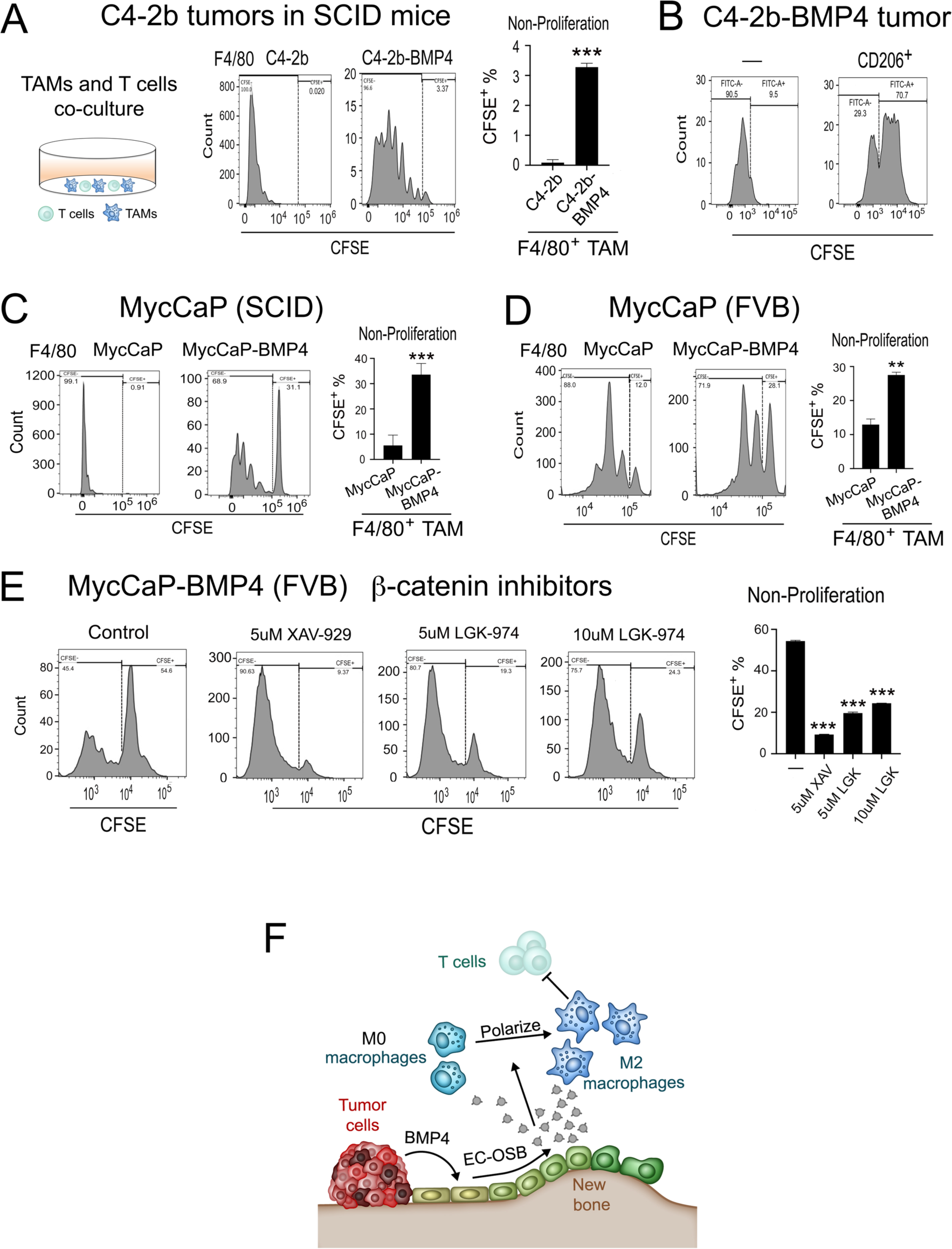
Bone-TAMs inhibit T cell proliferation through Wnt pathway. (A) CD45^+^CD11b^+^Gr1^−^F4/80^+^ macrophages from C4-2b-BMP4 tumors inhibit T cell proliferation. (B) CD45^+^CD11b^+^Gr1^−^F4/80^+^/CD206^+^ macrophages from C4-2b-BMP4 tumors inhibit T cell proliferation. CD206^+^ macrophages from C4-2b-vec tumors were too few to be included in the T cell proliferation assay. (C) CD45^+^CD11b^+^Gr1^−^F4/80^+^ macrophages from MycCaP-BMP4/SCID tumors showed higher inhibition of T-cell proliferation than those from MycCaP/SCID tumors. (D) CD45^+^CD11b^+^Gr1^−^F4/80^+^ macrophages from MycCaP-BMP4/FVB tumors showed higher inhibition of T-cell proliferation than those from MycCaP/FVB tumors. (E) Treatment of bone-TAMs from MycCaP-BMP4 tumors in FVB mice with Wnt pathway inhibitors XAV-939 or LGK-974 attenuate the effect of bone-TAMs on T-cell inhibition. (F) Graphical summary. EC-OSB cells-secreted factors promote M2 polarization of macrophages that inhibit T-cell proliferation. Error bars indicate SD. *p < 0.05, **p < 0.01, **p < 0.001, Student’s t test.

### Wnt pathway inhibition reduced bone-TAMs’ T-cell inhibitory activity

We next tested whether Wnt pathway activation in bone-TAMs was involved in their ability to suppress T-cell activity. Bone-TAMs isolated from MycCaP-BMP4 tumors generated in FVB mice were pre-treated with Wnt pathway inhibitors XAV-929 or LGK-974 for 1 hr before co-culture with T cells. XAV-929 or LGK-974 attenuated the inhibitory effect of bone-TAMs on T-cell proliferation (**Fig. 8E**). Similar results were observed with bone-TAMs isolated from MycCaP-BMP4 tumors generated in SCID mice (**Supplementary Fig. 9A**). These results suggest that Wnt pathway activation in bone-TAMs plays a role in T cell inhibition. To test whether XAV-929 or LGK-974 directly affected T cell proliferation, XAV-929 (5 uM) or LGK-974 (2.5, 5, 10 uM)-treated T cells were compared with vehicle-treated T cells, and found that they had similar proliferation rates (**Supplementary Fig. 9B**), suggesting that increased T-cell proliferation by XAV-929 or LGK-974 was unlikely to have resulted from a direct effect on T cells. Together, these results show that PCa-induced EC-OSB hybrid cell-secreted factors lead to M2 polarization of macrophages, which inhibit T-cell proliferation (**Fig. 8F**).

## Discussion

In this study, we reveal that osteoblastic bone metastases have an immunosuppressive bone-tumor microenvironment. We also mechanistically link the unique EC-to-OSB transition, which leads to PCa-induced osteoblastic bone metastases, as an “upstream” event that drives the “downstream” immunosuppression in the bone-TME (**Fig. 8F**). Our study thus unveils a novel mechanism that may account for the relative resistance of bone metastatic PCa to immune checkpoint therapy, and suggests therapeutic strategies that inhibit PCa-induced EC-to-OSB transition may reverse immunosuppression to promote immunotherapeutic strategies in bone metastatic prostate cancer. We have previously identified a novel therapy for targeting PCa-induced EC-to-OSB transition through a non-canonical retinoic acid receptor mechanism (20). Our study thus offers a strategy to improve immunotherapeutic responses.

Our study showed that CXCL14 is one of the EC-OSB-secreted factors that plays a role in the polarization and recruitment of M2-like macrophages to osteogenic tumors. Delineating CXCL14-mediated signaling pathways in macrophages may provide targets for reducing the immunosuppressive bone TME in PCa bone metastasis. Recent evidence suggested that CXCL14 receptor activation is a complex and cell type-dependent process (41,42). Chemokine receptors, including CXCR4 (43,44), GPR85 (45), and ACKR2 (46,47) have been implicated in CXCL14 function. CXCR4 is a member of the G protein-coupled receptor family and ERK1/2 is part of CXCR4 signal pathways (48). We also observed that CXCL14 induces ERK phosphorylation. Knockdown of CXCL14 in EC-OSB cells reduced the EC-OSB-mediated ERK phosphorylation in BMDMs and RAW264.7 cells (**Supplementary Fig. 10**). Furthermore, addition of recombinant mouse CXCL14 protein to BMDMs or RAW264.7 cells stimulated ERK phosphorylation (**Supplementary Fig, 10**). In ovarian cancer, CXCL14 was reported to activate β-catenin (40), however, we could not detect an increase in nuclear β-catenin by treating BMDMs with recombinant CXCL14 (**Supplementary Fig, 10**). Whether CXCL14-mediated polarization and recruitment of M2-like macrophages are through ERK1/2-pathway requires further investigation.

M1-like and M2-like macrophages are a continuum phenotype not easily characterized by single markers (31,49). Although we measured the total population of M2-like macrophages in our study, we recognize that this population contains many subtypes. Due to their plasticity, TAMs can switch from one type to another depending on soluble and cellular factors present in the tumor microenvironment (50). In addition to Wnt pathway, bone-TAMs may contain M2-like macrophages that are activated by other pathways. It is also possible that M2-like macrophages in different osteogenic tumors, e.g., MDA-PCa-118b, C4-2b-BMP4, MycCaP-BMP4, have different subtypes due to the influence of tumor secreted factors in addition to EC-OSB cells.

Macrophages are known to secrete various factors that regulate T-cell activity. Studies by Kfoury et al. (5) showed that in human PCa bone metastasis, macrophage-secreted CCL20 interacts with its receptor CCR6 on T cells to induce T-cell exhaustion. We did not find upregulation of CCL20 in the RNAseq analysis of bone-TAMs. However, we identified other secreted factors, including cytokines (e.g., IL11) (**Supplementary Table 2**), that are overexpressed in bone-TAMs. IL11 is implicated in immune regulation in colon cancer (51). Our RNAseq analysis also found that bone-TAMs have increased expression of Tph1 and Ido1, two key tryptophan metabolism enzymes implicated in immunosuppression (52,53). The mechanisms by which bone-TAMs regulate T-cell activities are currently being studied.

Given the significance of M2-like macrophages in TME and their dominant role in tumor growth and progression (54), strategies targeting immunosuppressive macrophages are being developed for cancer treatment (49,55,56). Among them, antibody or pharmacological agents targeting CSF1/CSF1R are being examined clinically for the treatment of various cancer types (57). Our studies showed that the properties of TAMs are continuously influenced by the recruitment and polarization signals from the tumor-induced bone reflecting an adaptive biological system (58). Since adaptive mechanisms of therapy resistance are driven by the tumor microenvironment (59), it is unlikely that targeting M2-like macrophages will be sufficient to reduce the immunosuppressive bone-TME. Our data suggest that inhibiting PCa-induced bone would improve immunotherapeutic strategies for bone metastatic PCa by targeting the biological process that drives the adaptive resistance. This was supported by our preclinical data showing that inhibition of the EC-to-OSB transition, via genetic, small molecules or Ra223, in osteogenic PCa tumors led to reduction of M2-like macrophages in the bone-TME. Clinical trials to test this hypothesis are in development.

## Materials and Methods

### Cell lines, antibodies, and reagents

Cell lines, antibodies, and reagents are listed in **Supplementary Table 3**. Human cell lines were authenticated using short tandem repeat. Cells were routinely tested to be mycoplasma free using MycoAlert Mycoplasma Detection Kit (LONZA, LT07-418). CD8^+^ T cells isolated from mouse spleen were cultured in RPMI1640 supplemented with 10% heat inactivation FBS and 100 U/mL penicillin/streptomycin.

### RNAseq data base analysis

To examine gene expression with M1/M2 macrophage markers in human prostate cancer, publicly available primary and mCRPC RNA-seq datasets (15) were downloaded from cBioportal. An M2-like tumor associated macrophage gene signature described previously (14) was used for hierarchical clustering of human bmCRPC samples. Patients with M2-high or M2-low gene signatures were compared for overall survival.

### Immunohistochemistry of human and mouse prostate cancer specimens

Formalin-fixed, paraffin-embedded human PCa specimens from primary tumors (4 cases), lymph node metastasis (4 cases), and bone metastasis (19 cases) were obtained from MDACC Prostate Cancer Tissue Bank through an institutional approved IRB protocol. Written informed consent were obtained before the study took place. Immunohistochemistry using CD68, F4/80 and CD206 antibodies on human PCa specimens or tumors generated from prostate cancer cells in SCID or FVB mice was performed using procedures described previously (17).

### Generation of C4-2b, C4-2b-BMP4, MycCaP, MycCaP-BMP4 tumors

Male SCID mice were randomly assigned to generate subcutaneous C4-2b or C4-2b-BMP4 tumors as described (20). Similarly, MycCaP and MycCaP-BMP4 tumors were generated in male SCID or FVB mice.

### Isolation of tumor-associated macrophages and flow cytometry

Subcutaneous tumors (∼ 1g) were cut into small pieces and digested with in 2.4 mL of dissociation solution [DMEM medium supplemented with enzyme mixture (100ul Enzyme D, 50uL Enzyme R and 12.5uL Enzyme A)] using Tumor Dissociation Kit (Miltenyi Biotec). After incubation for 45 minutes at 37°C, cells that passed through a 70-um cell strainer (BD Falcon) were collected as “1^st^ digested cells”. Bone containing tissues that were left on the 70-um cell strainer were further digested with 0.2% Collagenase II (Gibco) in PBS for 30 minutes at 37°C. Cells that passed through strainer were collected as “2^nd^ digested cells”. The 1^st^ and 2^nd^ digested cells were combined and treated with RBC lysis solution. The cells were washed with PBS and stained with LIVE/DEAD viability dyes, CD45, CD11b (Life Technologies), Gr-1 (BioLegend), F4/80 (BD Bioscience), and/or CD206 (BioLegend) mAb for 40 minutes at room temperature. Macrophage populations were analyzed and sorted by CytoFLEX SRT Cell Sorter (BECKMAN COULTER Life Science).

### Mouse primary BMDM isolation and M1 or M2 macrophage polarization

Bone marrow of femur and tibia of 8-12 weeks wild type C57B6 mice were flushed with DMEM medium and filtered through a 70 μM cell strainer. After treating with RBC lysis buffer, the cells in DMEM culture medium were plated onto a 10-cm tissue culture plate (1.5×10^7^/plate). After overnight incubation, supernatants were aspirated and replenished with fresh DMEM culture medium containing 15% L929 cell-conditioned medium. After refreshing medium at day 4, the cells were ready as primary macrophages (M0 macrophage) at day 7. To differentiate M0 macrophages to M1 or M2 macrophages, M0 macrophages were starved in 0.05% BSA DMEM medium overnight. The starved cells were then treated with 60 ng/mL LPS or 40 ng/mL IL4 for 72 hours to differentiate into M1 or M2 macrophages, respectively. A similar procedure was used to for differentiate of RAW or iBMDM cells to M1 or M2 macrophages, except the incubation time was 24 hours.

### Macrophage migration assay

For the M2 macrophage recruitment assay, cells (3 × 10^5^) in 300 mL of serum-free medium were seeded into FluoroBlock TM Cell Culture inserts (BD Falcon). Serum free media with 100 ng/mL LOX, 50 or 100ng/ml CXCL14, 2H11 control or EC-OSB conditional medium was placed in the lower chamber of a 24 well plate individually, using DMEM medium as control. After incubation for 14 hours, migrated cells were labeled with Calcein AM and were quantified in five randomly chosen visual fields.

### qRT-PCR

Total RNA was isolated using RNeasy Mini Kit (Qiagen). cDNA was prepared using cDNA Synthesis Kit (applied biosystems). qRT-PCR was performed as previously described (20) using mouse-specific primer sequences (**Supplementary Table 4**). mRNA levels were normalized to GAPDH and presented as relative mRNA expression or fold change.

### Immunoblotting and immunofluorescence

2H11 or EC-OSB cell lysates (20 μg) or conditional media (20 uL) were analyzed by immunoblotting with indicated antibodies. For nuclear fraction of control TAMs, bone TAMs, or bone-TAMs treated with Wnt inhibitor for 24 hours, TAMs were lysed in lysis buffer and the pellets were lysed in RIPA buffer to isolate nuclear fractions as previously described (17). For immunofluorescence analysis, cells were fixed with ice-cold 100% methanol and immunostained with the indicated antibodies as previously described (17,20).

### Knockdown of LOX or CXCL14 in 2H11 cells

MISSION pLKO.1 lentiviral shRNA (MilliPORE Sigma) was used to knockdown LOX or CXCL14 in 2H11 cells. The shRNA clones were selected with 5 µg/mL puromycin. Cells infected with empty pLKO.1 lentiviral vector were used as controls. shRNA sequences are listed in **Supplementary Table 4**.

### Mineralization assays

2H11 cells, 2H11-shLOX-#1,#5 or 2H11-shCXCL14-#2,4,5 were cultured in serum-free DMEM medium overnight followed by treatment with or without BMP4 (100 ng/mL) for 48 h, then switched to osteoblast differentiation medium (α-MEM containing 10%FBS, 100 μg/ml ascorbic acid and 5 mmol/L β-glycerophosphate) for 14 days with fresh medium changes every three days as previously described (20).

### Generation of MDA PCa-118b tumors treat with ATRA, Palovarotene, LDN or Rad223

MDA PCa-118b (1×10^6^) were injected into SCID mice subcutaneously. After 3 weeks, mice bearing MDA PCa-118b tumors with similar tumor size were treated with ATRA, Palo, or LDN as previously described (20). Treatment of C4-2b-BMP4 tumors with Rad223 was as previously described (60).

### Generation of MycCaP-BMP4 cell line by dilution cloning

cDNA for mouse BMP4 was first cloned into FUW-Luc-mCh-puro vector. The Luciferase, mCherry and Puro markers were then excised from the plasmid to generate FUW-BMP4 vector. FUW-BMP4 vector was transfected into MycCaP cells and diluted into 96-well plates. The expression of BMP4 protein in the culture supernatants was detected by Western blot. Clones that expressed higher levels of BMP4 were further diluted and screened. Clones that expressed relatively high levels of BMP4 were selected as MycCaP-BMP4 cell line.

### RNA-seq of tumors and tumor-associated macrophages

For isolating RNAs for RNAseq analysis of MycCaP or MycCaP/BMP4 tumors, MycCaP or MycCaP/BMP4 tumors were cut to small pieces, homogenized in RLT lysis buffer (RNAeasy Mini Kit (Qiagen)) using economical bench top homogenizer (PT1600E), and RNA isolated using the RNAeasy Mini Kit (Qiagen). For RNAseq analysis of tumor-associated macrophage, the digested cells from MycCaP or MycCaP-BMP4 tumor were FACS sorted for CD45^+^, CD11b^+^, GR1^−^ and F4/80^+^ cells and RNAs were prepared. RNA-seq analysis was performed at University of Texas Health Cancer Genomics Center (Houston, TX).

### CD8+ T cell isolation and T cell suppression assay

Mouse spleen (∼ 0.1g) from 8-12-week-old C57B6 mice was placed in C-Tube (Miltenyi Biotec). After mechanical disruption, the cells that passed through 70-uM cell strainer were purified with EasySep^TM^ Mouse CD8^+^ T Cell Isolation Kit (STEMCELL Technologies Inc). For T-cell proliferation assays, T cells were labeled with CFSE, stimulated with CD3/CD28 T-Activator and mouse IL-2 (Gibco), and cocultured with F4/80^+^ macrophages at 2:1 (T cells:macrophages) ratios in 96-well flat-bottom plates. Seventy-two hours later, cells were analyzed by flow cytometry. For suppression assay, Bone-TAMs isolated from MycCaP-BMP4 tumors generated in FVB mice were pre-treated with XAV-929 or LGK-974 for 1 hour before co-cultured with T cells.

### Study approval

Animal studies have been approved by the Institutional Animal Care and Use Committee at M.D. Anderson Cancer Center.

### Statistical analysis

Data were expressed as the mean ± S.D. p < 0.05 by Student’s *t*-test was considered statistically significant.

## Data availability

All data reported in this paper will be shared by the lead contact upon request.

## Author contributions

G.Y., P.G.C., G.W., S.-H.L. conceived the idea, planned the experiments, and wrote the manuscript. G.Y., C.Z.L.M., X.L., J.H.S., S.-C.L. carried out the experiments. G.Y., M.Z., P.T., X.S., J.L., J.Z., M.P.M., G.W., S.-H.L. performed data analysis and interpretation. T.P., C.J.L., G.W., S.-H.L. provided scientific inputs for the development of the project. All the co-authors critically reviewed the present manuscript before submission.

