## Supplemental figures for "Prostate cancer-induced endothelial-to-osteoblast transition generates an immunosuppressive bone tumor microenvironment"

Supplementary Figure 1

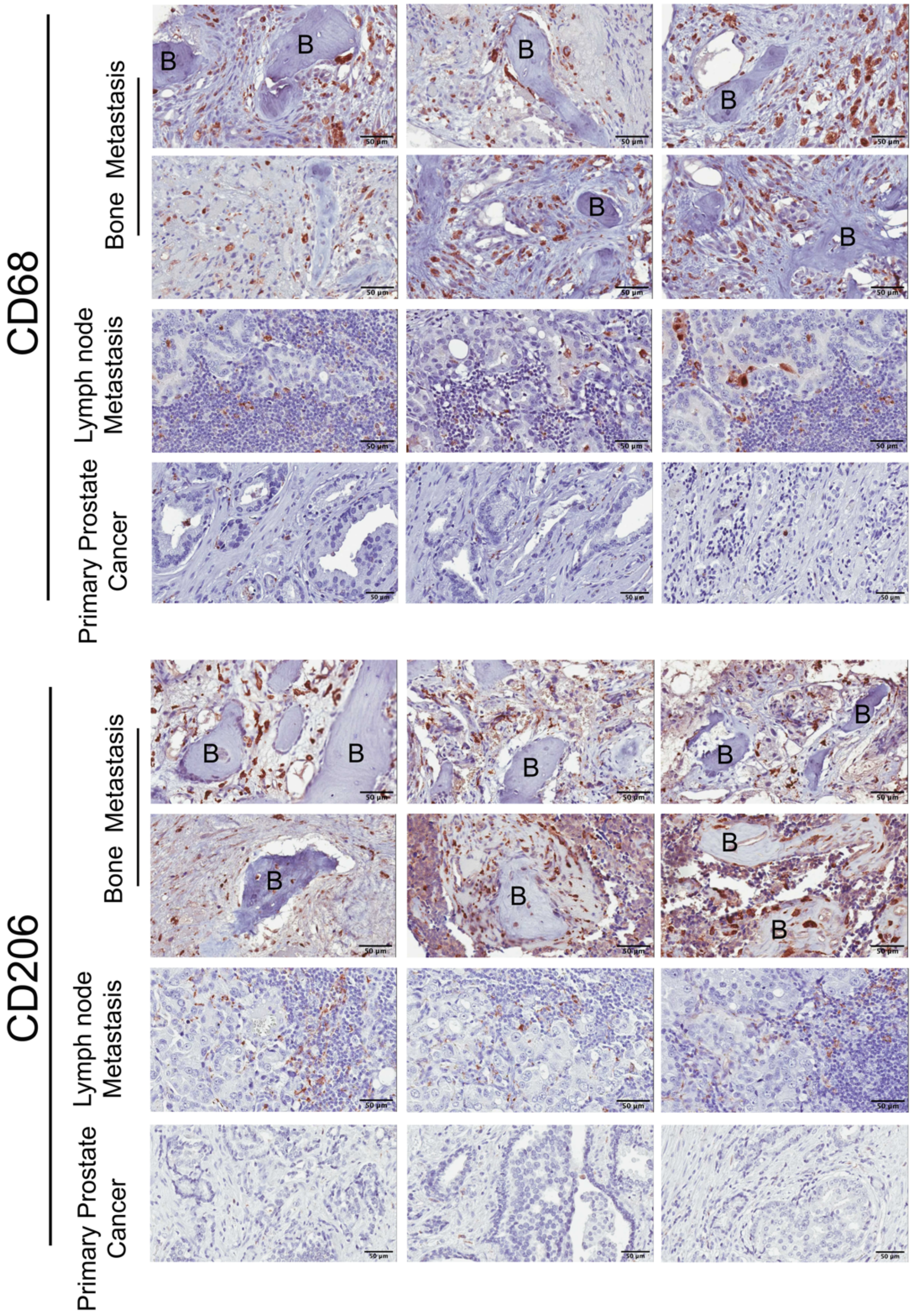

**Supplementary Figure 1. Immunohistochemistry of CD68⁺ and CD206⁺ cells in specimens from bone metastasis, lymph node metastasis, or primary prostate cancer.**

Supplementary Figure 2

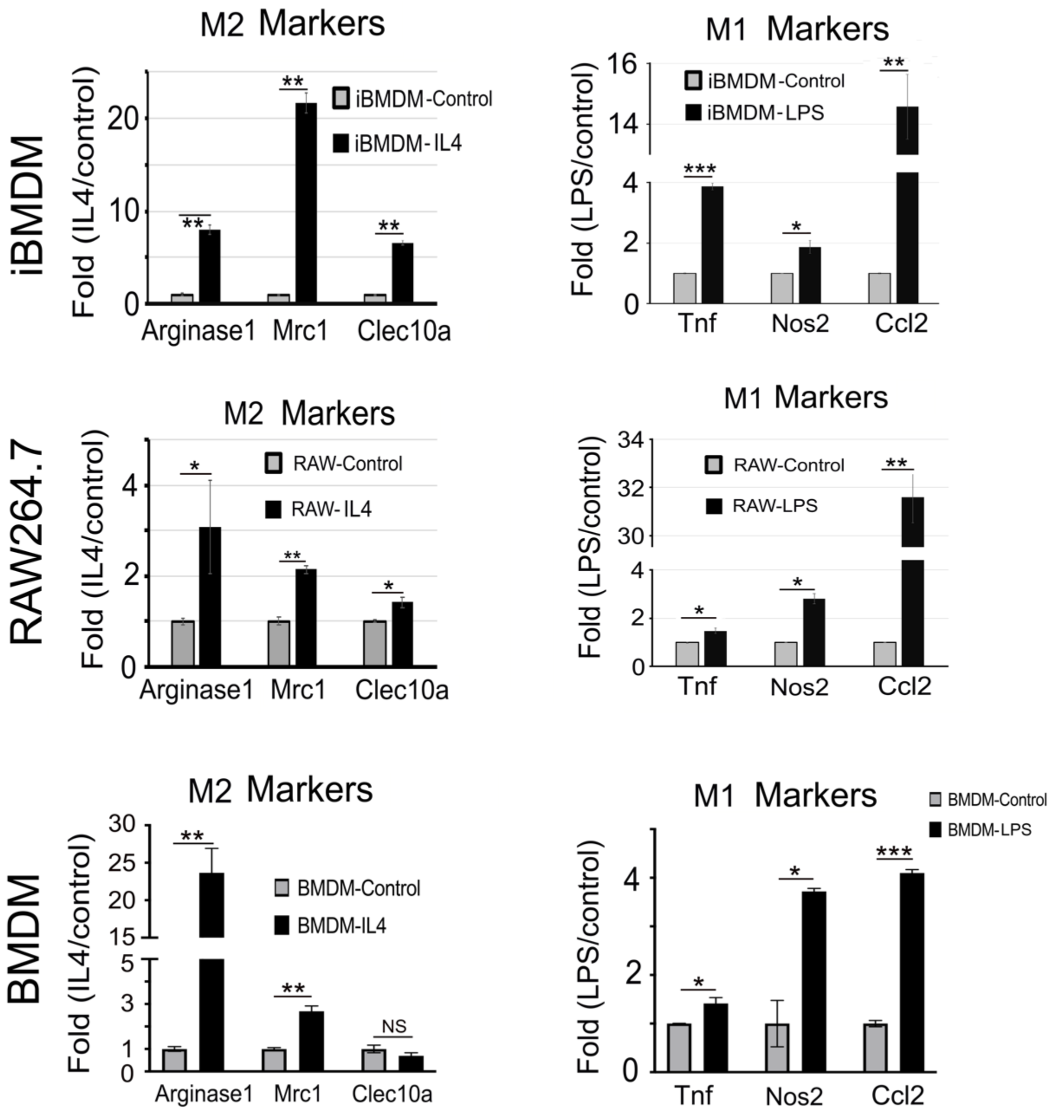

**Supplementary Figure 2.** iBMDM and RAW264.7 cells were induced to polarize to M2-like or M1 macrophage by incubating with IL-4 or lipopolysaccharide (LPS), respectively, for 24 hr. qRT-PCR showed the upregulation of M2 macrophage markers *arginase1 (Arg1), Mrc1(CD206)* and *Clec10a* by IL-4, and the upregulation of M1 macrophage markers *Tnf, Nos2,* and *Ccl2.* Polarization of BMDMs was performed similarly, except that BMDMs were incubated with IL-4 or LPS for 72 hrs.

### Supplementary Figure 3

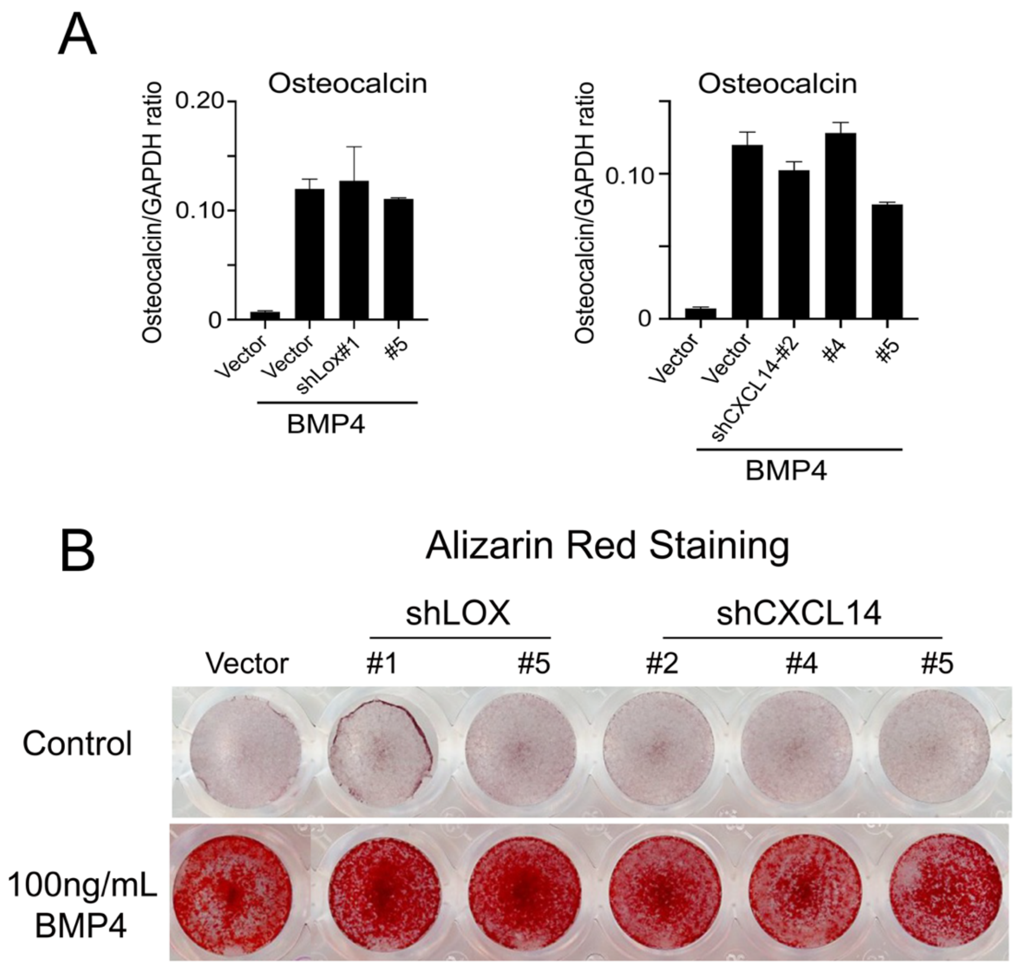

**Supplementary Figure 3. Knockdown of *Lox* or *Cxcl14* in 2H11 cells did not have significant effects on BMP4-mediated EC-to-OSB transition.** shRNAs were used to knock down *Lox or Cxcl14* in 2H11 endothelial cells. The cells were treated with BMP4 (100 ng/mL) for 48 hours to induce EC-to-OSB transition. qRT-PCR was used to determine the expression of *osteocalcin* and *osterix*, factors upregulated in osteoblasts. For osteoblast differentiation, BMP4-treated cells were further cultured in osteoblast differentiation medium for 14 days with medium change every 3 days and mineralization was determined by Alizarin staining. Knockdown of *Lox* or *Cxcl14* in 2H11 cells did not have significant effects on BMP4-mediated EC-to-OSB transition.

Supplementary Figure 4

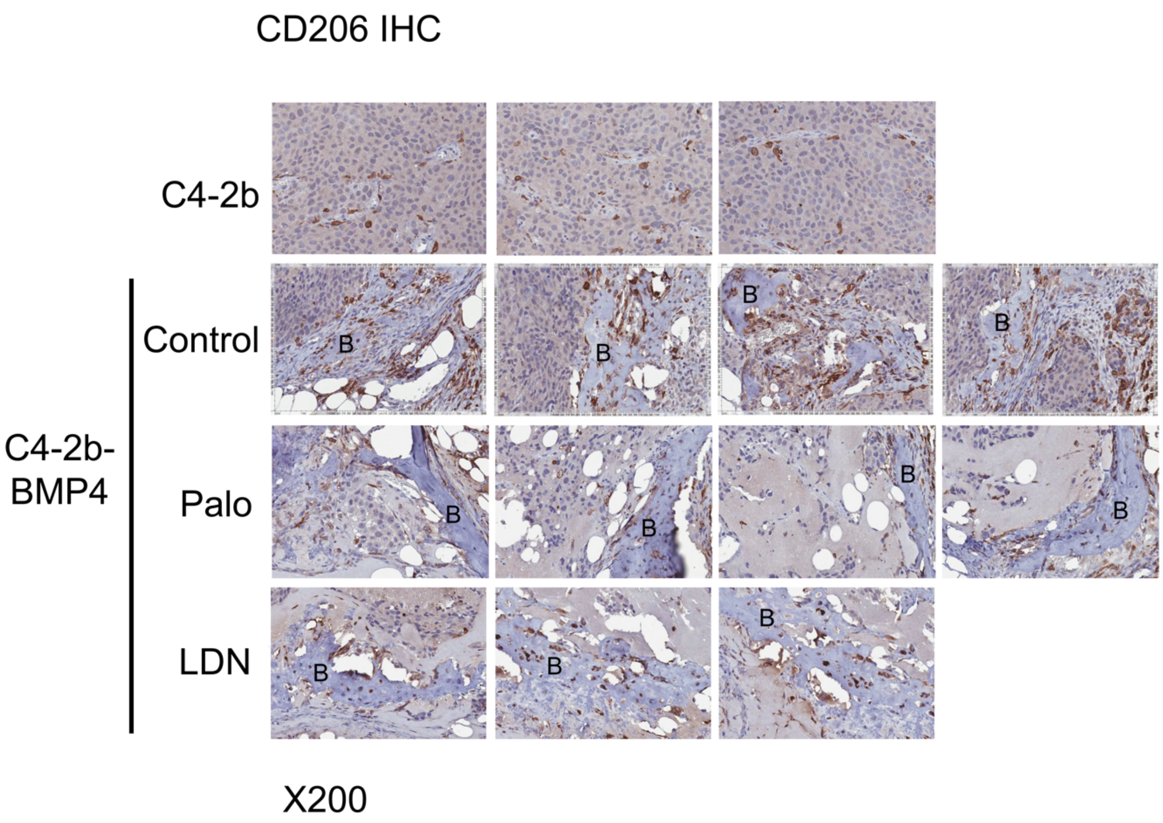

**Supplemental Fig. 4. Palovarotene or LDN193189 reduced CD206$^+$ cells in C4-2b-BMP4 tumors.** IHC of CD206 in C4-2b-BMP4 tumors from mice treated with LDN193189 or palovarotene. The levels of CD206$^+$ macrophages in LDN193189 or palovarotene-treated tumors were reduced compared to control C4-2b-BMP4 tumors.

### Supplementary Figure 5

Palo and LDN on macrophage proliferation

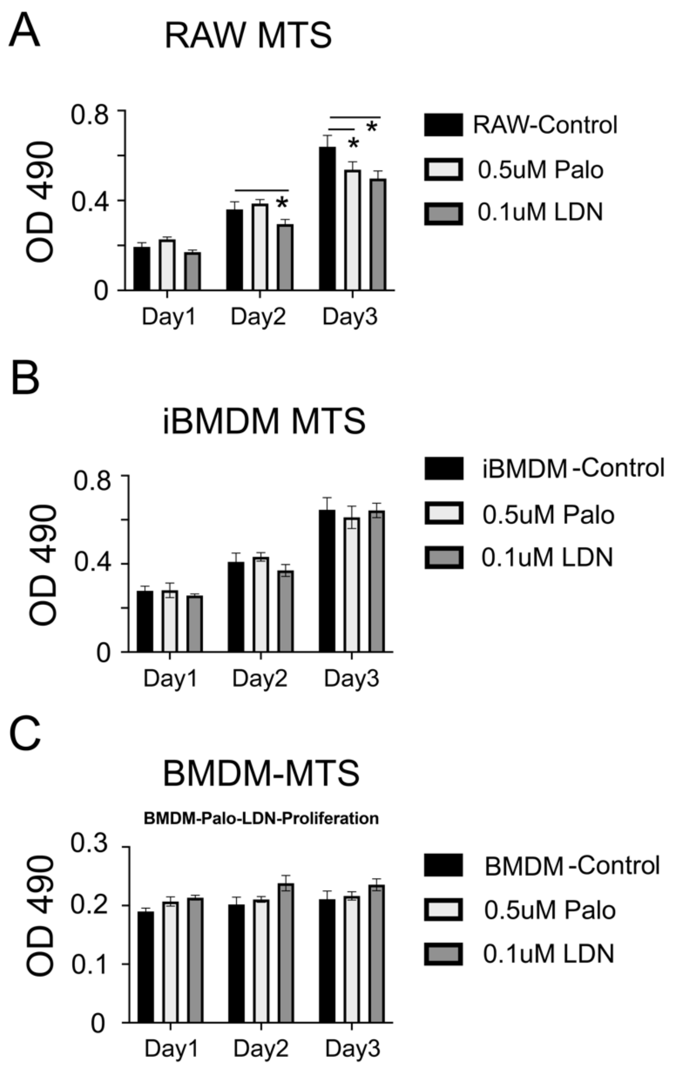

**Supplementary Fig. 5. Palovarotene and LDN193189 do not affect macrophages proliferation *in vitro*.** Macrophages in culture were treated with palovarotene (0.5 uM) or LDN193189 (0.1 uM) for 1-3 days. MTS assay was used to determine cell proliferation. *p<0.05

#### Supplementary Figure 6

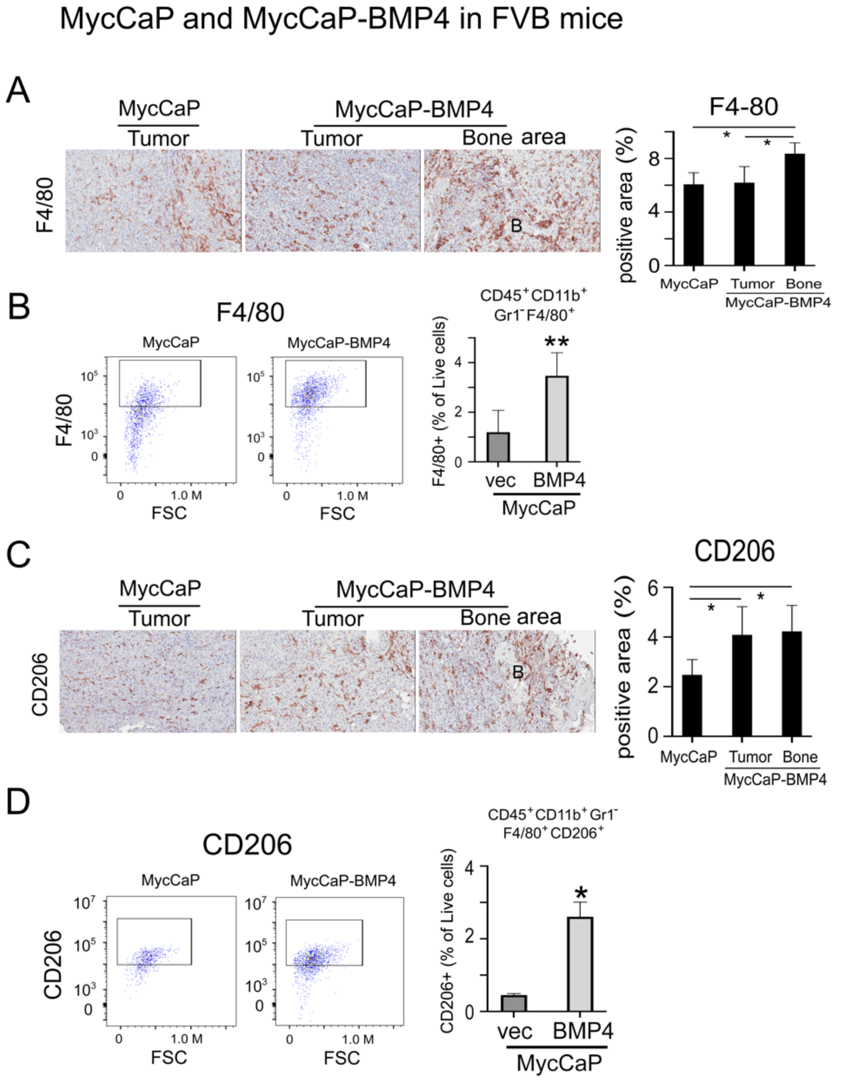

**Supplementary Figure 6. MycCaP-BMP4 tumors in FVB mice showed recruitment of F4/80$^{+}$ and CD206$^{+}$ macrophages to the tumor-induced bone area.**

(A) F4/80 IHC of MycCaP and MycCaP-BMP4 tumors generated in FVB mice. (B) FACS analysis of CD45$^{+}$CD11b$^{+}$Gr$^{-}$F4/80$^{+}$ cells from MycCaP and MycCaP-BMP4 tumors generated in FVB mice. (C) CD206 IHC of MycCaP and MycCaP-BMP4 tumors generated in FVB mice. (D) FACS analysis of CD45$^{+}$CD11b$^{+}$Gr$^{-}$CD206$^{+}$ cells from MycCaP and MycCaP-BMP4 tumors generated in FVB mice. F4/80$^{+}$ and CD206$^{+}$ macrophages were enriched in MycCaP-BMP4 tumors in the area with tumor-induced bone. The levels of CD45$^{+}$CD11b$^{+}$Gr1$^{-}$F4/80$^{+}$ and CD45$^{+}$CD11b$^{+}$Gr1$^{-}$F4/80$^{+}$CD206$^{+}$ in MycCaP-BMP4 tumors were higher compared to those in MycCaP tumors.

Supplementary Figure 7

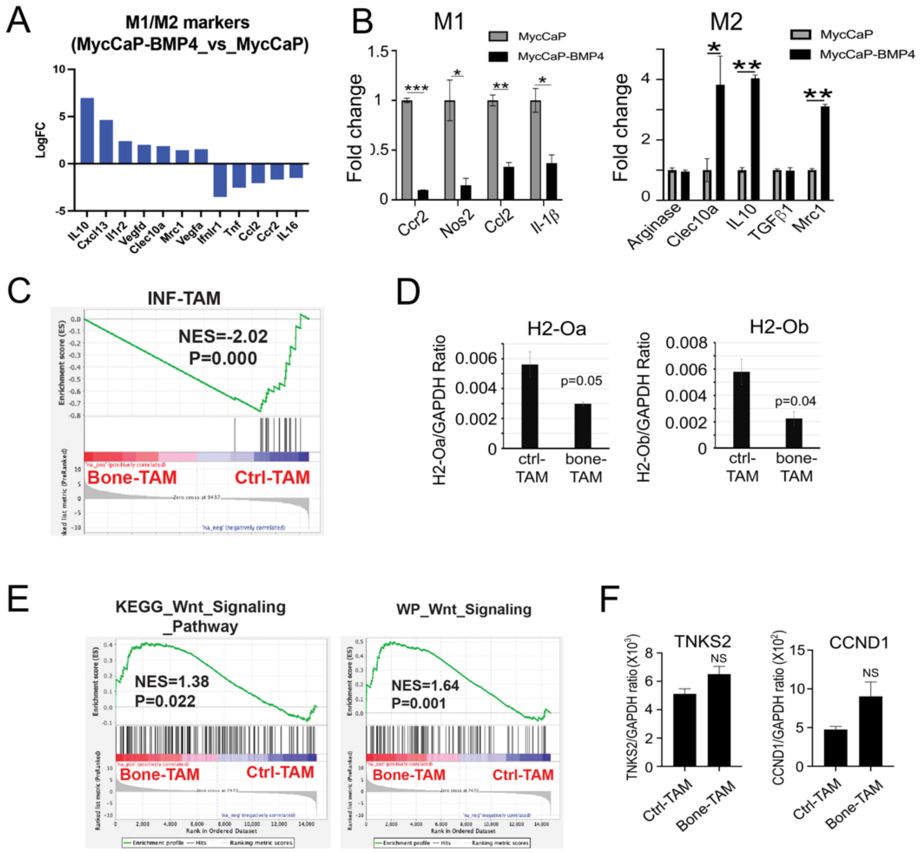

**Supplementary Figure 7. RNAseq analysis of tumor-associated macrophages showed enrichment of M2 macrophage-related genes and upregulation of Wnt signaling pathways in MycCaP-BMP4 compared to MycCaP tumors.**

(A) Upregulation of M2 markers and downregulation of M1 markers in MycCaP-BMP4 compared MycCaP tumors.

(B) qRT-PCR of M1-macrophage and M2-macrophage markers in total RNAs isolated from MycCaP and MycCaP-BMP4 tumors. MycCaP tumors expressed higher levels of M1-macrophage markers, while MycCaP-BMP4 tumors expressed higher levels of M2-macrophage markers.

(C) GSEA showed that IFN-TAM gene signature was downregulated in bone-TAM.

(D) The expression levels of MHC-II genes H2-Oa and H2-Ob were lower in in bone-TAMs compared to ctrl-TAMs.

(E) GSEA analysis showed Wnt signaling is activated in bone-TAMs compared to ctrl-TAMs.

(F) qRT-PCR of the mRNA levels of TNKS2 and CCND1. These two downstream canonical Wnt pathway target genes were not significantly upregulated in bone-TAMs compared to ctrl-TAM.

### Supplementary Figure 8

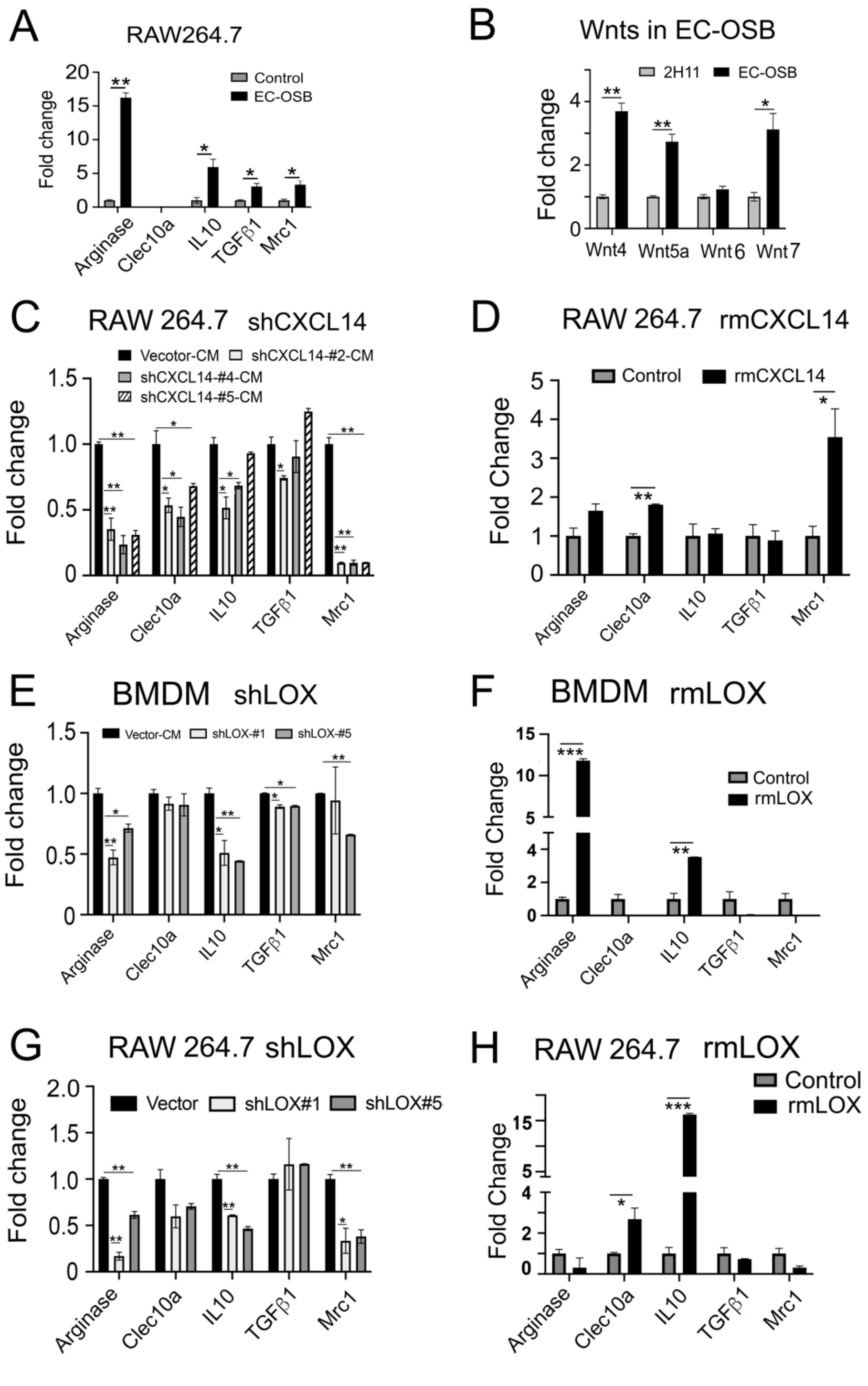

**Supplementary Figure 8. Factors secreted from EC-OSB cells promote M2 polarization of macrophages.**

(A) Treatment of RAW264.7 with EC-OSB CM upregulated markers of M2-like macrophages.

(B) EC-OSB hybrid cells secrete several Wnt ligands as detected by qRT-PCR.

(C) Treatment of RAW264.7 with conditioned media from EC-OSB cells with knockdown of *CXCL14* (sh*Cxcl14*) reduced the M2 marker expression.

(D) Treatment of RAW264.7 cells with recombinant mouse CXCL14 upregulated M2 marker expression.

(E) Conditioned media from EC-OSB cells with knockdown of *Lox* (sh*Lox*) reduced the M2 markers of BMDM compared with shvector EC-OSB CM.

(F) Treatment of BMDMs with recombinant mouse LOX protein upregulated M2 markers.

(G) Treatment of RAW264.7 with conditioned media from EC-OSB cells with knockdown of *Lox* (sh*Lox*) reduced the M2 marker expression.

(H) Treatment of RAW264.7 cells with recombinant mouse LOX upregulated M2 marker expression.

Error bars indicate SD. *$p < 0.05$, **$p < 0.01$, **$p < 0.001$, Student's t test.

### Supplementary Figure 9

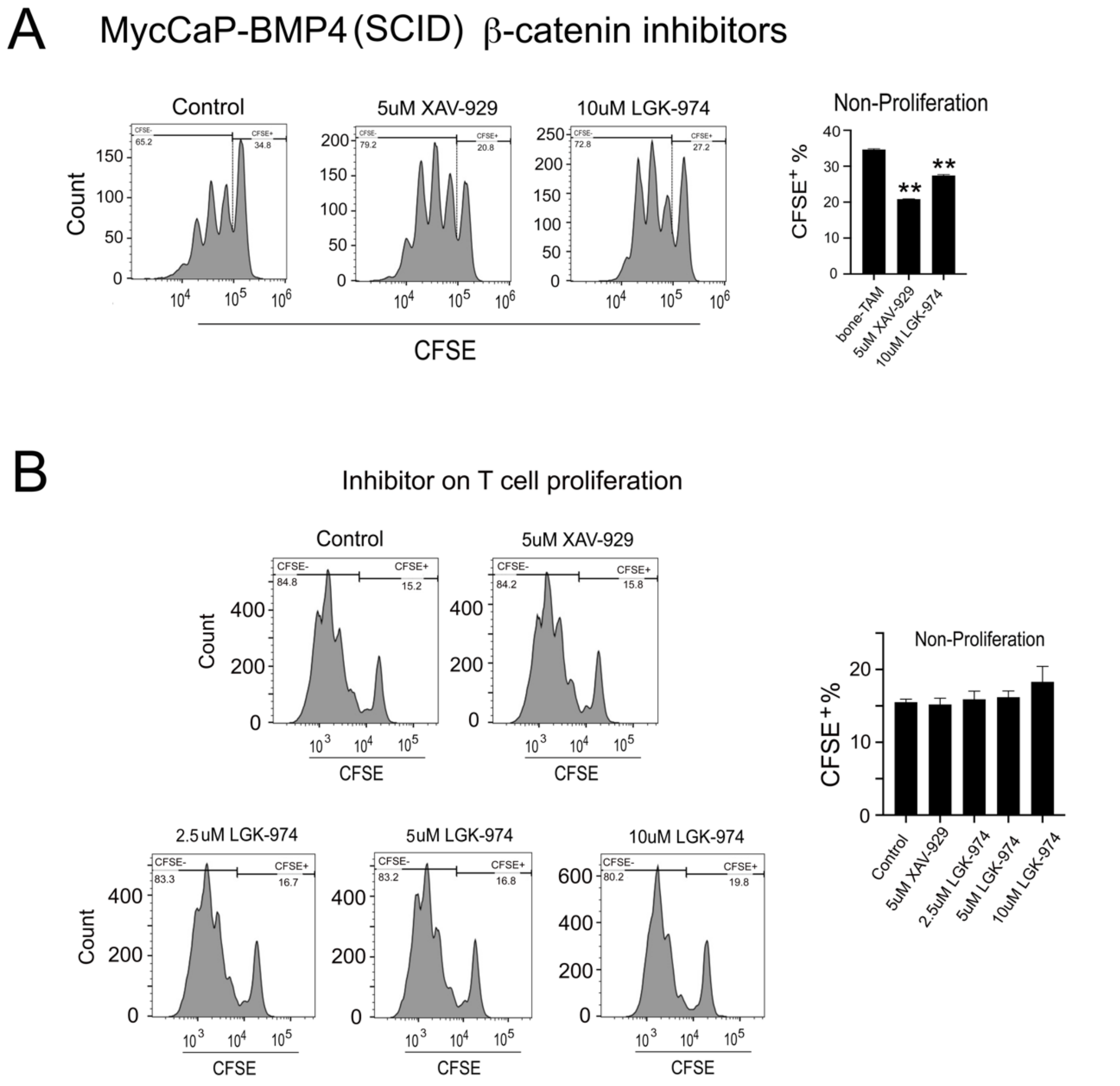

**Supplementary Figure 9. Bone-TAMs inhibit T cell proliferation through Wnt pathway.**

(A) Treatment of bone-TAMs from MycCaP-BBMP4 tumors in SCID mice with Wnt pathway inhibitors XAV-939 or LGK-974 attenuate the effect of bone-TAMs on T-cell inhibition.

(B) XAV-929 or LGK-974 showed no significant effects on T cell proliferation. CFSE-labeled T cells were treated with XAV-929 at 5 uM or LGK-974 at 2.5, 5, or 10 uM for 72 hr. FACS analysis showed no significant effect on T cell proliferation at these concentrations.

Supplementary Figure 10

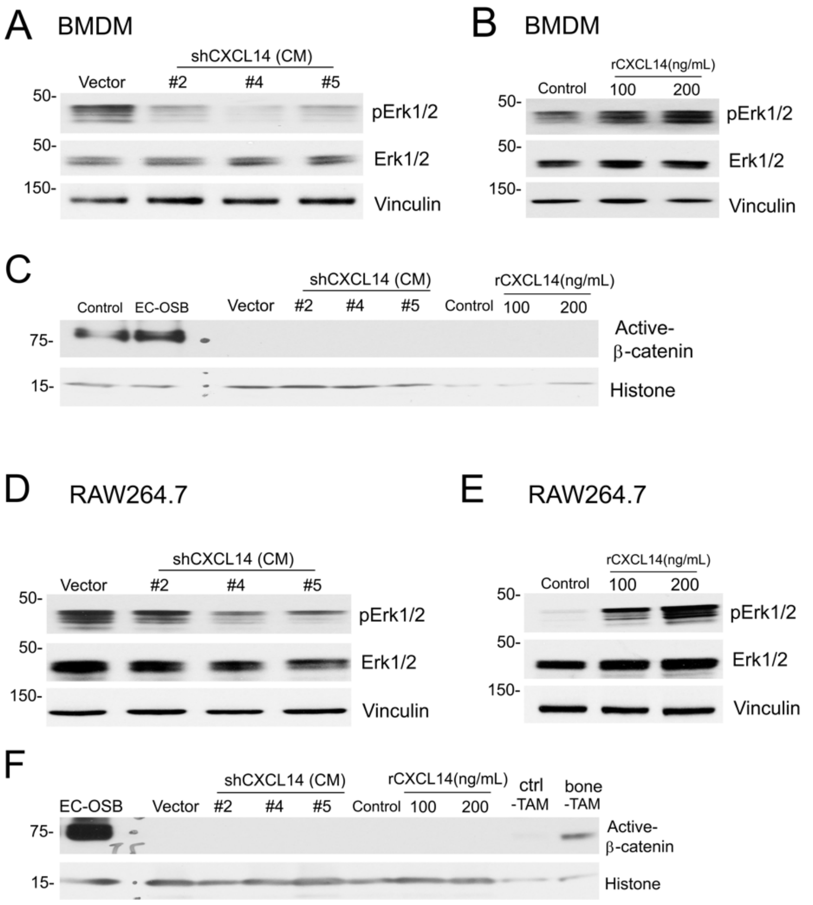

**Supplementary Figure 10. CXCL14 induces ERK phosphorylation in BMDMs and RAW264.7 cells.** Conditioned media from EC-OSB cells were used to treat BMDMs or RAW264.7 cells. (A) Knockdown of CXCL14 in EC-OSB cells reduced the EC-OSB-mediated ERK phosphorylation in BMDMs. (B) Addition of recombinant mouse CXCL14 protein to BMDMs increased ERK phosphorylation. (C) Active β-catenin was not detected in the nuclear fraction of control, shCXCL14 EC-OSB cell conditioned media-treated, or recombinant mouse CXCL14 protein-treated BMDMs. Nuclear fraction of control (untreated 2H11 cells) and EC-OSB cells (BMP4-treated 2H11 cells) were used as positive controls. (D) Knockdown of CXCL14 in EC-OSB cells reduced the EC-OSB-mediated ERK phosphorylation in RAW264.7 cells. (E) Addition of recombinant mouse CXCL14 protein to RAW264.7 cells increased ERK phosphorylation. (F) Active β-catenin was not detected in the nuclear fraction of control, shCXCL14 EC-OSB cell conditioned media-treated, or recombinant mouse CXCL14 protein-treated RAW264.7 cells. Nuclear fraction of EC-OSB cells, ctrl-TAMs, and bone-TAMs were used as controls.

**Supplementary Table 3. List of reagents.**

| Reagent | Source | Catalogue Number |
|---|---|---|
| **Antibodies** | | |
| CD68, macrosialin | Dako | M0814, RRID: AB_2314148 |
| hMMR MAb (Cl 685645) | R&D Systems | MAB25341,RRID:AB_10890782 |
| F4/80(D2S9R) XP Rabbit mAb | Cell Signaling Technology | 70076, RRID:AB_2799771 |
| mMMR Aff Pur PAb | R&D Systems | AF2535, RRID:AB_2063012 |
| Recombinant Anti-LOX antibody | Abcam | Ab174316, RRID:AB_2630343 |
| CXCL14 antibody | Abcam | Ab137541, RRID: |
| Non-phospho(Active)β-Catenis(Ser45) | Cell Signaling Technology | 19807, RRID:AB_2650576 |
| h/mWnt-5a MAb (Cl 442625) | R&D Systems | MAB645- SP, RRID: 10571221 |
| Phosphp-GSK-3β | Cell Signaling Technology | 9336,RRID:AB_331405 |
| GSK-3β(27C10) | Cell Signaling Technology | 9315,RRID:AB_490890 |
| ROR1(D6T8C) | Cell Signaling Technology | 16540,RRID:AB_2798764 |
| p-CaMKIIα (Thr 286) | Santa Cruz Biotechnology | sc-12886-R, AB_2067915 |
| CAMK2 | Proteintech | 13730-1-AP, RRID:AB_2070320 |
| GAPDH(D16H11) | Cell Signaling Technology | 5174, RRID:AB_10622025 |
| Vinculin(E1E9V)XP | Cell Signaling Technology | 13901,RRID:AB_2728768 |
| LaminA/C | Cell Signaling Technology | 2032S; RRID:AB_2136278 |
| eBioscience Fixable Viability Dye eFluor 506 | Life Technologies | 65- 0866-14, RRID: |
| CD45 Monoclonal Antibody (30-F11), eFluor 450, eBioscience | Life Technologies | 48- 0451-82, RRID:AB_1518806 |
| CD11b Monoclonal Antibody (M1/70), APCeFluor 780, eBioscience | Life Technologies | 47- 0112-82,RRID:AB_1603193 |
| APC anti-mouse Ly-6G/Ly-6C(Gr-1) Antibody | BioLegend | 198411; RRID:AB_313376 |
| Ms F4/80 PE-CF594 T45-2342 | BD Bioscience | 565613; RRID:AB_2734770 |
| PerCP/Cyanine5.5 anti-mouse CD206 (MMR) [Clone: C068C2] | BioLegend | 141715; RRID: AB_2561991 |
| **Inhibitors** | | |
| XAV-939 | Selleckchem | S1180 |
| Vismodegib(GDC-0449) | Selleckchem | S1082 |
| LDN 193189 | Axon Medchem | Axon 1509 |

| | | |
|---|---|---|
| Palovarotene | Toronto Research Chemicals | P165900 |
| ATRA | Millipore-Sigma | S1574 |
| **Recombinant Protein** | | |
| Recombinant human BMP4 | R&D Systems | 314-BP |
| Lysyl Oxidase (LOX) Recombinant, Mouse | United States Biological | 155736 |
| Recombinant mouse CXCL14/BRAK | R&D Systems, Inc. | 730-XC-025 |
| rh/mWnt-5a | R&D Systems, Inc. | 645-WN-010 |
| **Cell Lines** | | |
| 2H11 mouse lymphoid endothelial cells | ATCC | CRL-2163, RRID:CVCL_6762 |
| RAW264.7 | ATCC | TIB-71™, RRID:CVCL_0493 |
| MycCaP | ATCC | # CRL-3255, RRID:CVCL_J703 |
| C4-2b | ATCC | #CRL-3315 |
| NCTC CLONE 929 AREOLAR FIBROBL | ATCC | #CCL-1 RRID:CVCL_0462 |
| **Kit** | | |
| Mouse CXCL14/BRAK ELISA Kit | Novus Biologicals Inc | # NBP2-70015 |
| EasySepTrademark Mouse CD45 Positive Selection Ki | STEMCELL Technologies Inc | #18945 |
| EasySep™Mouse $CD8^+$ Cell Isolation Kit | STEMCELL Technologies Inc | #19853 |
| Tumor Dissociation Kit | Miltenyi Biotec | #130-096-730 |

**Supplementary Table 4. Oligonucleotide sequences for qRT-PCR and shRNA knockdown.**

| Name | Sequence (5'-3') | product length(bp) |
|---|---|---|
| LOX-qPCR-F | TCTTCTGCTGCGTGACAACC | |
| LOX-qPCR-R | GAGAAACCAGCTTGGAACCAG | 117 |
| CXCL14-qPCR-F | GGCCCAAGATCCGCTACA | |
| CXCL14-qPCR-R | TGGGTACTTTGGCTTCATTTCC | 56 |
| Osteocalcin-qPCR-F | GCTCTGTCTCTCTGACCTCA | |
| Osteocalcin-qPCR-R | TGGACATGAAGGCTTTGTCA | 68 |
| CCR2-qPCR-F | TCCTTGGGAATGAGTAACTGTGT | |
| CCR2-qPCR-R | TGGAGAGATACCTTCGGAACTT | 142 |
| Nos2-qPCR-F | GCCCAGCCAGCCCAAC | |
| Nos2-qPCR-R | TGATGGACCCCAAGCAAGAC | 240 |
| CCl2-qPCR-F | AGCTCTCTCTTCCTCCACCA | |
| CCl2-qPCR-R | GGTCAGCACAGACCTCTCTC | 590 |
| IL-1B-qPCR-F | ACCCCAAAAGATGAAGGGCTG | |
| IL-1B-qPCR-R | TACTGCCTGCCTGAAGCTCT | 112 |
| IL6-qPCR-F | CCCCAATTTCCAATGCTCTCC | |
| IL6-qPCR-R | CGCACTAGGTTTGCCGAGTA | 141 |
| TNFa-qPCR-F | GGCAGTTAGGCATGGGATGA | |
| TNFa-qPCR-R | TACCTACGACGTGGGCTACA | 186 |
| Arginase 1-qPCR-F | CGTGAGTTGCAGTTCTGCTC | |
| Arginase 1-qPCR-R | CTTGGCCCAGCACCGTATAA | 722 |
| Clec10a-qPCR-F | TGGTGGTCGTCTCCGTGATT | |
| Clec10a-qPCR-R | ACAGATCCAGCGGAAGGTTCTC | 746 |
| IL10-qPCR-F | GGCCCAGAAATCAAGGAGCA | |
| IL10-qPCR-R | CACACTGCAGGTGTTTTAGCTT | 249 |
| TGFβ1-qPCR-F | CATCCATGACATGAACCGGC | |
| TGFβ1-qPCR-R | GAAGTTGGCATGGTAGCCCT | 217 |
| Mrc1-qPCR-F | GTCAGAACAGACTGCGTGGA | |
| Mrc1-qPCR-R | AGGGATCGCCTGTTTTCCAG | 281 |
| Cxcl9-qPCR-F | GGAGTTCGAGGAACCCTAGTG | |
| Cxcl9-qPCR-R | GGGATTTGTAGTGGATCGTGC | 82 |
| Cxcl10-qPCR-F | CCAAGTGCTGCCGTCATTTTC | |
| Cxcl10-qPCR-R | GGCTCGCAGGGATGATTTCAA | 157 |
| Cxcl11-qPCR-F | GGCTTCCTTATGTTCAAACAGGG | |
| Cxcl11-qPCR-R | GCCGTTACTCGGGTAAATTACA | 108 |
| H2-Oa-qPCR-F | TCTACCAATCTTACGACGCTTCT | |
| H2-Oa-qPCR-R | CACACGACCTCCTCGTTCT | 94 |
| H2-Ob-qPCR-F | AGGCGGACTGTTACTTCACC | |
| H2-Ob-qPCR-R | ATCCAGGCGTTTGTTCCACTG | 164 |
| TNKS1-qPCR-F | CCCAGGACCCTCACTCCAT | |
| TNKS1-qPCR-R | TCCCAAACTCCCAGTCTTGAA | 117 |
| TNKS2-qPCR-F | CGCCCGAGAAGGTGAACAG | |
| TNKS2-qPCR-R | TTTGCACCGTTCTGAAGAAGAT | 115 |
| CCND1-qPCR-F | GGGCAGCCCCAACAACTTCC | |
| CCND1-qPCR-R | TCCTCAGTGGCCTTGGGGTC | 234 |
| MET-qPCR-F | GTGAACATGAAGTATCAGCTCCC | |
| MET-qPCR-R | TGTAGTTTGTGGCTCCGAGAT | 100 |
| Myc-qPCR-F | ATGCCCCTCAACGTGAACTTC | |
| Myc-qPCR-R | CGCAACATAGGATGGAGAGC | 228 |
| Ror1-qPCR-F | TGAGCCGATGAATAACATCACAA | |
| Ror1-qPCR-R | CAGGTGCATCATTCTTGAACCA | 110 |
| Wnt7b-qPCR-F | TTTGGCGTCCTCTACGTGAAG | |

| | | |
|---|---|---|
| Wnt7b-qPCR-R | CCCCGATCACAATGATGGCA | 145 |
| Wnt5a-qPCR-F | CAACTGGCAGGACTTTCTCAA | |
| Wnt5a-qPCR-R | CCTTCTCCAATGTACTGCATGTG | 77 |
| Wnt4-qPCR-F | AGACGTGCGAGAAACTCAAAG | |
| Wnt4-qPCR-R | GGAACTGGTATTGGCACTCCT | 126 |
| Wnt6-qPCR-F | GCAAGACTGGGGGTTCGAG | |
| Wnt6-qPCR-R | CCTGACAACCACACTGTAGGAG | 202 |
| FZD3-qPCR-F | ATGGCTGTGAGCTGGATTGTC | |
| FZD3-qPCR-R | GGCACATCCTCAAGGTTATAGGT | 109 |
| DVL1-qPCR-F | GCTGACTGTGAAGAGTGA | |
| DVL1-qPCR-R | GCATTGGCAATGGTGAT | 108 |
| Vangl2-qPCR-F | ACTCGGGCTATTCCTACAAGT | |
| Vangl2-qPCR-R | TGATTTATCTCCACGACTCCCAT | 110 |
| GAPDH-qPCR-F | TGCAGTGGCAAAGTGGAGAT | |
| GAPDH-qPCR-R | TTTGCCGTGAGTGGAGTATA | 96 |
| mBMP4-BamH1-F | GGATCCATGATTCCTGGTAACCGAATGCTG | |
| mBMP4-EcoR-H-R | GAATTCTCATCAGCGGCATCCACACCCCTCT | 1224 |

#### shRNA

| Gene | Accessions | Clone ID | Mature Antisense (5'-3') | Target |
|---|---|---|---|---|
| Lox | NM_010728 | TRCN0000011850 (#1) | GCTGCACAATTTCACCGTATT | CDS |
| | NM_010728 | TRCN0000011849 (#5) | CCTGGATGTTATGACACCTAT | CDS |
| CXCL14 | NM_019568 | TRCN0000065369 (#2) | CTGCGAGGAGAAGATGGTTAT | CDS |
| | NM_019568 | TRCN0000065371 (#4) | ACGGGTCCAAGTGTAAGTGTT | CDS |
| | NM_019568 | TRCN0000065372 (#5) | GCTGGAAATGAAGCCAAAGTA | CDS |
